# Versatile labeling and detection of endogenous proteins using tag-assisted split enzyme complementation

**DOI:** 10.1101/2020.12.01.407072

**Authors:** Suraj Makhija, David Brown, Struan Bourke, Yina Wang, Shuqin Zhou, Rachel Rudlaff, Rasmi Cheloor-Kovilakam, Bo Huang

## Abstract

Recent advances in genome engineering have expanded our capabilities to study proteins in their natural states. In particular, the ease and scalability of knocking-in small peptide tags has enabled high throughput tagging and analysis of endogenous proteins. To improve enrichment capacities and expand the functionality of knock-ins using short tags, we developed the tag-assisted split enzyme complementation (TASEC) approach, which uses two orthogonal small peptide tags and their cognate binders to conditionally drive complementation of a split enzyme upon labeled protein expression. Using this approach, we have engineered and optimized the tag-assisted split HaloTag complementation system (TA-splitHalo) and demonstrated its versatile applications in improving the efficiency of knock-in cell enrichment, detection of protein-protein interaction, and isolation of biallelic gene edited cells through multiplexing.

## Introduction

CRISPR/Cas9-mediated genome engineering techniques have revolutionized the study of endogenous biology. With these techniques, one powerful application is to label proteins by genomic knock-in so that the abundance, dynamics, and interactions of endogenous proteins can be examined while avoiding artifacts of overexpression. For this purpose, one approach is to use fluorescent protein (FP) fusions, enabling the use of fluorescence activated cell sorting (FACS) to directly isolate and enrich for knocked-in (KI) cells. However, the large size of FPs leads to potential perturbation of the tagged protein’s localization and function and more importantly, impacts the efficiency and scalability of the knock-in approach.

In contrast, short peptide tags can be used to overcome these limitations, but they are not inherently fluorescent and are not compatible with live cell FACS unless the tag is extracellularly localized and therefore compatible with antibody staining. An alternative option is split fluorescent protein, or FP_11_ tags, which were developed based on the self-complementing split GFP_1-10/11_ ^1,2^ and the split of mNeonGreen ^3^ and sfCherry ^4^. These tags are 16 a.a. peptides derived from the 11^th^ β strand of FPs. Once expressed, the corresponding FP_1-10_ fragment will bind FP_11_ tags to form a functional FP. Owing to their combined small size and fluorescence, FP_11_ tags have facilitated the generation and analysis of mammalian cell libraries containing endogenously tagged proteins^5^.

Still, FP_11_ tags have intrinsic limitations in fluorophore brightness and photostability, making it challenging to detect and track low expression targets. Moreover, it is highly desirable to expand this tagging approach to other split protein complementation systems, such as split luciferase for bioluminescence detection ^6^, split protease for synthetic circuits ^7^, and split enzymatic tags, particularly split HaloTag ^8^, that enable labeling of the target protein with organic fluorophores that are bright, photostable and available in many different colors. This also would enable reporter outputs beyond fluorescence. Unfortunately, none of these split proteins are self-complementing, meaning that they require additional protein-recruitment strategies to induce the complementation of the split fragments. In addition, the roughly central position of their split points means that neither fragment is small enough to serve as a short peptide tag. Therefore, they cannot be directly adapted to endogenous protein tagging like the split FP_1-10/11_ systems.

Here, we present a general approach that enables short peptide tagging of proteins to activate split protein complementation, which we named <u>t</u>ag <u>a</u>ssisted <u>s</u>plit <u>e</u>nzyme <u>c</u>omplementation (TASEC). For our model system, we focused on HaloTag, a self-labelling enzyme engineered to covalently bind chloroalkane ligands. This property makes HaloTag extremely versatile as available ligands for HaloTag include a range of “turn-on” fluorescent dyes with distinct spectral properties ^9^ and dyes optimized for single molecule tracking ^10^, super-resolution microscopy ^11^, and expansion microscopy ^12^. Based on an existing non-self-complementing split HaloTag ^8^, we have engineered the tag-assisted split HaloTag (TA-splitHalo) that utilizes two orthogonal, short peptide tags and their respective binders in living cells to scaffold the complementation of HaloTag on the target protein (Figure 1A). We have demonstrated the versatility of this system in the detection of low expression protein targets, the sorting of biallelic KI cells, and the detection of endogenous protein-protein interactions (Figure 1B).

**Figure 1:**
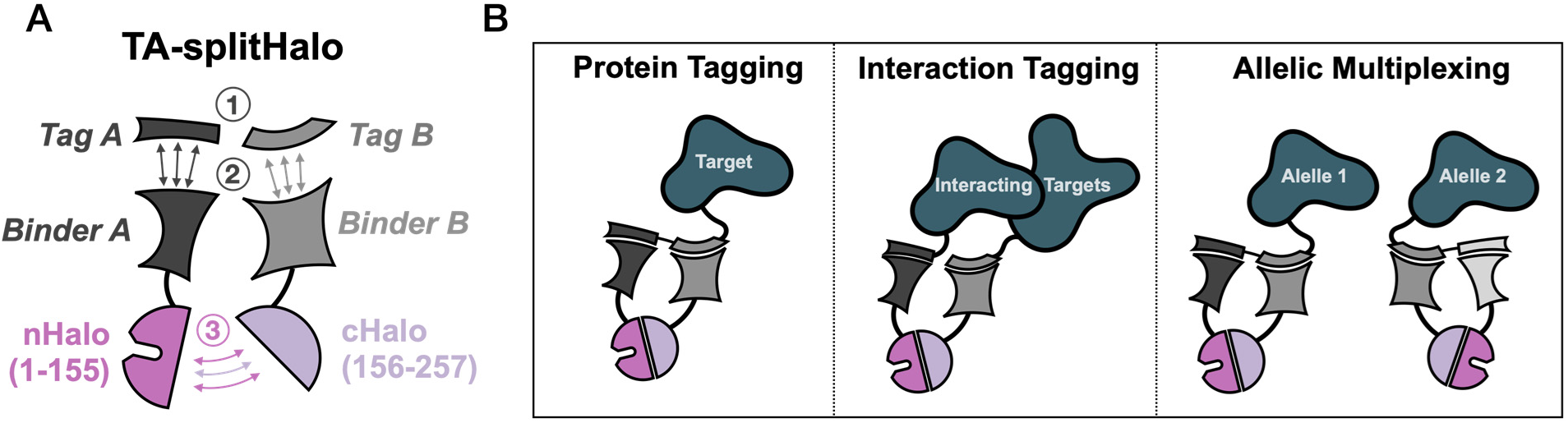
TA-splitHalo Overview and Applications. (A) A schematic of the TASEC concept as applied to TA-splitHalo. (1) Two orthogonal peptide tags arefused to the target protein(s). (2) Cognate binders fused to the two unfolded splitHalo fragments are recruited to the tags. (3) Confinement of the splitHalo fragments drives refolding of a functional HaloTag molecule. (B) The TA-splitHalo strategy can be applied to tag proteins by knocking-in both tags on the same target protein (left), protein interactions by knocking-in individual tags on interacting proteins (center), or tagging multiple alleles by assigning different TA-splitHalo approaches to different alleles in the same cell (right).

## Results and Discussion

### The engineering of TA-splitHalo systems

A TA-splitHalo ‘system’ consists of two orthogonal peptide tags and their respective binders, arranged in a way to drive efficient complementation of split HaloTag. To identify the set of tags and binders for TA-splitHalo systems and optimal architectures of, we employed a flow cytometry screening assay to test various combinations and arrangements.

The first system we tested in this manner was the GFP/Spy system. In this case, the tags were GFP_11_ and SpyTag002 (SpyT) and the respective binders were GFP_1-10_ and SpyCatcher002 (SpyC). There are 8 possible TA-splitHalo ‘architectures’ with the HaloTag fragments positioned at the N- or C-terminus of the two peptide binders. In the GFP/Spy case, we named these architectures GS01 to GS08. Our numerical nomenclature is standardized for all splitHalo architectures where the SpyC and positioning of splitHalo components are the same for each numbered construct. Structural representations of all architectures and these equivalencies are shown in Figure S1.

In an ideal architecture, the TA-splitHalo fragments should only fold if the detector components are expressed and bound to a tagged target. We cloned all 8 possible detector architectures into a common “landing pad” backbone to create a split-Halo detection plasmid library. Since both fusion proteins are expressed on the same plasmid backbone with the same promoter, we can assume the same range of splitHalo fusion protein expression levels relative to one another. Additionally, this vector gave us the ability to generate single-copy cell lines of optimal architectures for subsequent studies.

To rank the architectures, we used GFP_11_-SpyT-mCherry as the bait, with mCherry giving a readout of tag expression. SpyT-mCherry was used as the negative control for the complementation specificity. In all experiments we used a JF646 HaloTag Ligand (referred to as JF646 throughout). The far-red, fluorogenic, JF646 probe avoids sources of cellular autofluorescence and therefore maximize the signal to background ratio.

We tested the GFP/Spy system by transfecting each detection plasmid alongside bait plasmids expressing either SpyT-mCherry or GFP_11_-SpyT-mCherry (Figure 2A). This was performed in an equimolar ratio with an equal number of cells per sample to minimize expression level variability. We developed Python tools to uniformly select singlet cell events and subsequently analyze the relationship between mCherry expression and reconstitution-derived splitHalo (Figure S2). From the raw data (Figure 2B), we obtained hit rates (Halo+ / mCherry+) for each architecture (Figure 2C). Welch’s unequal variances *t*-test of biological triplicates shows that all GFP/Spy architectures impart statistically significant (P < 0.05) conditionality for the condition with both tags as hypothesized (Figure S2). We picked architectures with the highest hit rates GS02 (P = 0.0006) and GS07 (P = 0.02) for further characterization. GS02 has greatest fold difference between the background hit rate and the true hit rate. Conversely, GS07 yields the highest splitHalo signal when both tags are present, but it also has the second most background of any GFP/Spy architecture.

**Figure 2:**
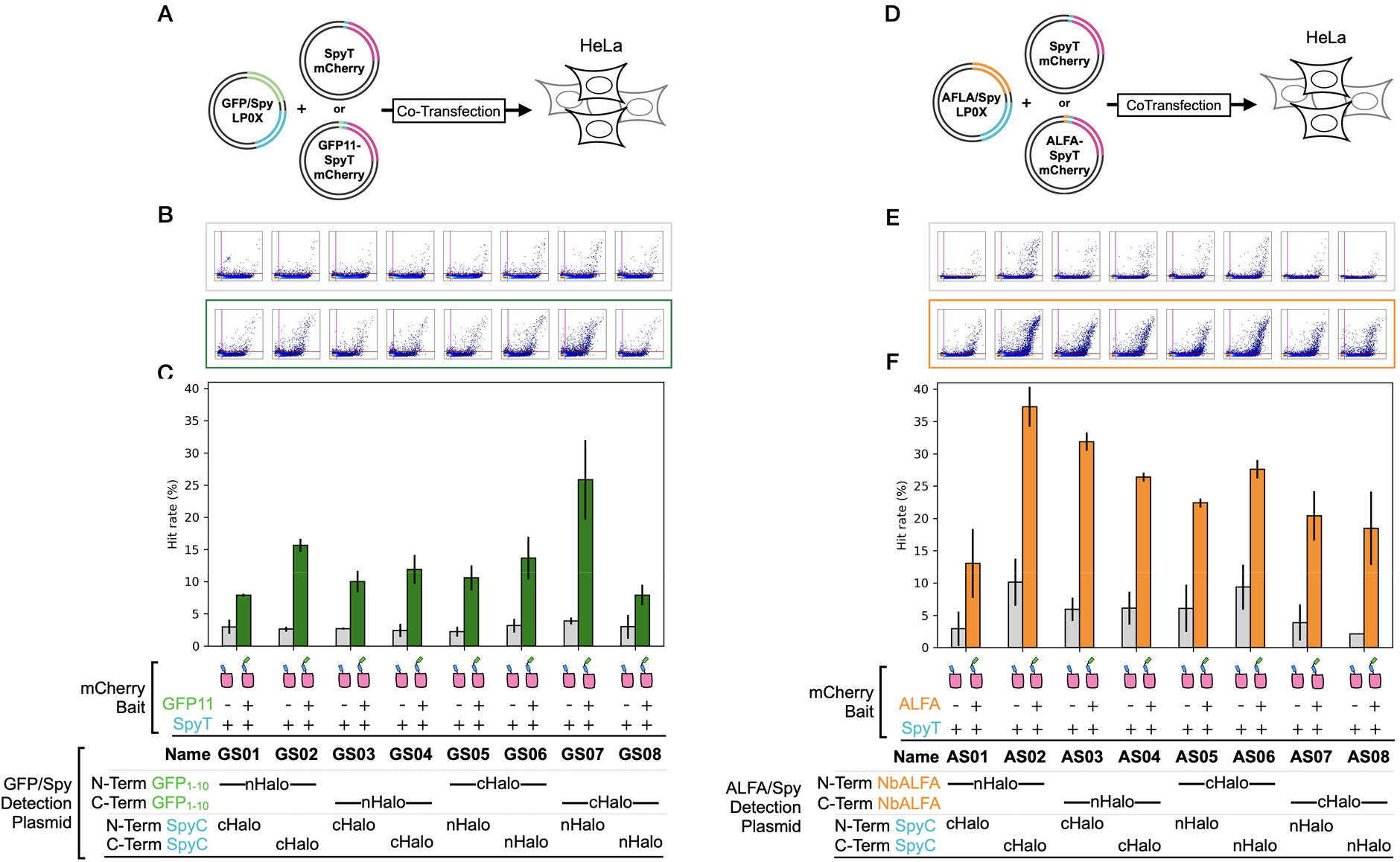
TA-splitHalo Architecture Scanning. (A) Schematic of GFP/Spy co-transfection architecture scan. Cells were transfected with a plasmid that expresses each GFP/Spy TA-splitHalo architecture and an mCherry bait expression vector tagged with SpyT alone or GFP11-SpyT. (B) Raw flow cytometry depicting GFP/Spy TA-splitHalo signal (y-axis) vs. mCherry tag reporter expression (x-axis). Each plot is a random sampling of 10k singlet-gated events for each architecture with a SpyT-mCherry bait (grey) or GFP11-SpyT-mCherry (green). (C) Mean (n=3) hit rate of each GFP/Spy splitHalo architecture in samples with SpyT-mCherry (grey) or GFP11-SpyT-mCherry (green). An equivalent analysis was done for ALFA/Spy architectures (D-F). For all architectures except AS01, differences from SpyT-mCherry were significant (P < 0.05) by Welch’s t-test (Figure S2).

The next system we tested was ALFA/Spy splitHalo. The ALFA tag is a structured α-helix peptide with a cognate nanobody named NbALFA that we employed as the binder ^13^. We held SpyT constant to make direct comparisons between various splitHalo systems. The ALFA/Spy system is dark until the addition of a Halo ligand because there are no extraneous fluorophores in the architectures.

We employed the same screening strategy to test the ALFA/Spy architectures, AS01 to AS08. In this case, we transfected each detection plasmid alongside bait plasmids expressing either SpyT-mCherry or ALFA-SpyT-mCherry in an equimolar ratio (Figure 2D). Again, the raw data (Figure 2E) were analyzed, and architecture-specific hit rates were obtained (Figure 2F). Like the GFP/Spy system, all architectures exhibited with statistically significant signal increases with two tags aside from AS01 (P = 0.06). From the ALFA/Spy system, we selected AS02 (P = 0.0007) and AS04 (P = 0.003) for further study. AS02 has the highest hit rate of the ALFA/Spy architectures while AS04 is another architecture that performed well with the SpyC-cHalo component. This allowed us to attribute the differences seen when comparing GS02, AS02, and AS04 solely to the varied nHalo fusion.

To further investigate the specificity of our four best performing architectures, we repeated our assay, adding an untagged mCherry bait to determine whether splitHalo background in the SpyT controls was the result of SpyT recruitment (Figure S2). Welch’s unequal variances *t*-test indicated that the slightly increased hit rate in co-transfections with SpyT-mCherry rather than untagged mCherry was not statistically significant. This means that any background we see is likely from highly transfected cells (Figure S2). In this independent biological experiment significant differences between SpyT-mCherry, and tandem tag-mCherry were again observed for GS07, AS02 and AS04, but not for GS02.

### Detection of knock-In cells using TA-splitHalo

To determine the utility of splitHalo systems for detecting successful KI events, we generated stable HEK293T cell lines to allow fair comparison between our selected split-Halo architectures and against the legacy split GFP_1-10/11_ platform. Via BxbI-driven integration, we created cell lines with single-copy integrants of the four selected split-Halo detection architectures (Figure S4). For comparison, we also created a GFP_1-10_ cell line in a similar manner. By placing the detection modules at the same genomic site - the AAVS1 safe harbor locus - we can compare the split-Halo systems to the split-GFP with the detection proteins present in the cell lines in known relative quantities. This way, we can compare KIs to the same target across all the cell lines and KI strategies.

After generating (Figure S4 and validating (Figure S5) the landing pad cell lines through genomic PCR we performed a KI targeting the LMNA gene in each cell line with the tagging strategy that corresponds with each detection module (Figure 3A-B). Using these sorted KIs, we characterized the KI populations against the original isogenic lines as controls comparing split-Halo systems and architectures to one another and to GFP_1-10/11_ (Figure 3C and D). Kolmogorov– Smirnov tests indicate highly significant (P < 0.001) increases in GFP signal for GFP_(1-10)_, GS02 and GS07 upon knock in. Increases in AS02 and AS04 were also highly significant but with much smaller effect sizes (0.04 and 0.05 respectively compared to 0.58 for GFP_(1-10)_, reflecting the absence of GFP in these cell lines (Figure S6). No significant increase in Halo signal was observed in the GFP_1-10_ cell line upon GFP_11_ knock in (P = 0.11) as expected for a control lacking any HaloTag components. All four TA-splitHalo systems showed a highly significant increase in Halo signal upon knock-in (Figure S6).

**Figure 3:**
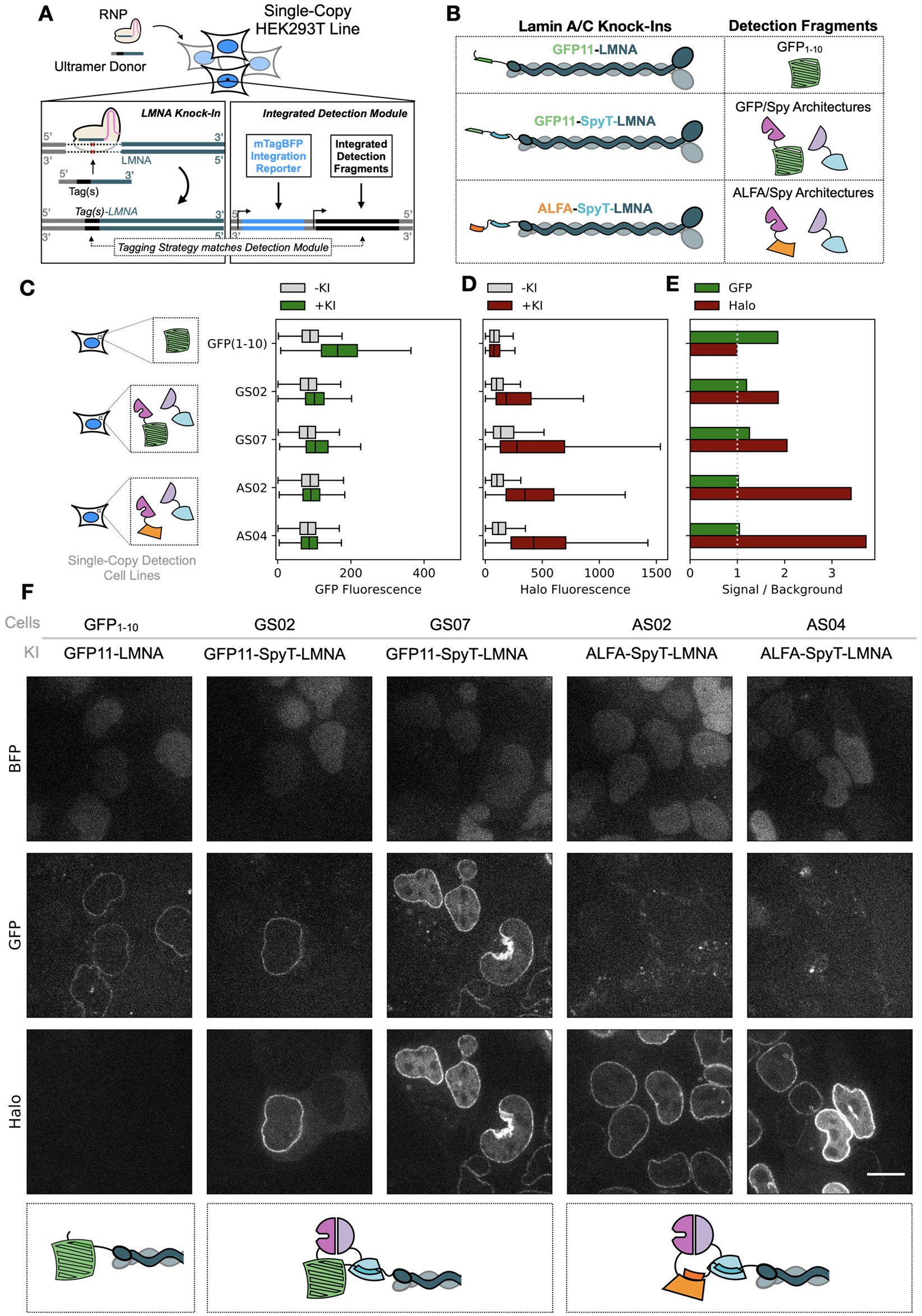
Evaluating Signal to Background of TA-splitHalo Architectures in Single-Copy Cell Lines. (A) Overview of knock-in strategy in single-copy detection cell lines. Short tag knock-ins on the LMNA gene were performed in cell lines pre-engineered to express the requisite detection components for each detection system off a single transcriptional unit at the same genomic locus. (B) Table depicts illustrations of the relevant proteins expressed in knock-in lines. (C) Median signal intensity in the GFP channel in background (grey) and knock-in (green) conditions in each detection cell line. (D) Median signal intensity in the TA-splitHalo channel in background (grey) and knock-in (red) conditions in each detection cell line. Median is derived from flow cytometry data of 10k cells per condition. (E) Signal to background values calculated by taking the ratio of knock-in to background GFP median signal (green) and TA-splitHalo signal (red) for each detection system. Dashed line depicts 1:1 signal to background detection threshold. (F) Confocal images of all LMNA knock-ins in detection cell line. Panels show nuclear BFP integration reporter (top row) LMNA-specific splitGFP signal (center row), and LMNA-specific TA-splitHalo signal (bottom row)

By taking the ratio of the median signal intensities for cell populations of the original cell lines and sorted LMNA KI lines in the GFP and splitHalo channels (Figure 3C-3D), we can compare signal to background ratios in each channel for the same KI target across detection platform (Figure 3E). The GFP_1-10/11_ system yields a 1.85 signal to background ratio in the landing pad cell line (Figure S6). In comparison, each of the TA-splitHalo architectures perform comparably or better in the far-red channel using 10 nM JF646 dye. In the ALFA/Spy architectures, the signal to background ratio was 3.4 and 3.7 for AS02 and AS04 respectively, demonstrating that splitHalo outperforms GFP_1-10/11_ for detection of LMNA KI.

Confocal imaging confirmed that both GFP and splitHalo signal had a nuclear envelope localization corresponding to lamin (Figure 3F). In the splitHalo systems, most of the background originates from basal levels of tag-independent splitHalo complementation. For architectures with higher median background like GS07, we see that this corresponds to visible cytoplasmic TA-splitHalo signal verifying that this unwanted signal is not driven by any single tag and is the result of non-specific complementation.

### Detecting protein-protein interactions with TA-splitHalo

After quantifying the performance of the strategy for knock-in detection, we sought to test whether TA-splitHalo could enrich for cells containing a protein-protein interaction. Because TA-splitHalo tagging systems consist of two peptide tags, we tested whether separating the tags and placing them on interacting proteins could yield a sortable signal upon complex formation or multimerization.

For this purpose, we used the homodimerization of lamin A/C chains as a model system, which places the N-termini of separate monomers in proximity^14,15^. We expected to see complemented Halo signal when the two tags of TA-splitHalo are present on different alleles of the LMNA gene.

Specifically, we modified our LMNA KI protocol to include two ultramer donor strands in the AS04 cell line, so that we can achieve simultaneous double-KI of ALFA-LMNA and SpyT-LMNA (Figure 4A) leading to TA-splitHalo complementation at lamin dimers (Figure 4B). Once we perform the KI and stain with JF646, Halo+ cells should contain both edits (Figure 4C). We enriched for this Halo+ population (Figure 4D) and confirmed nuclear envelope labelling using widefield imaging (Figure 4E).

**Figure 4.**
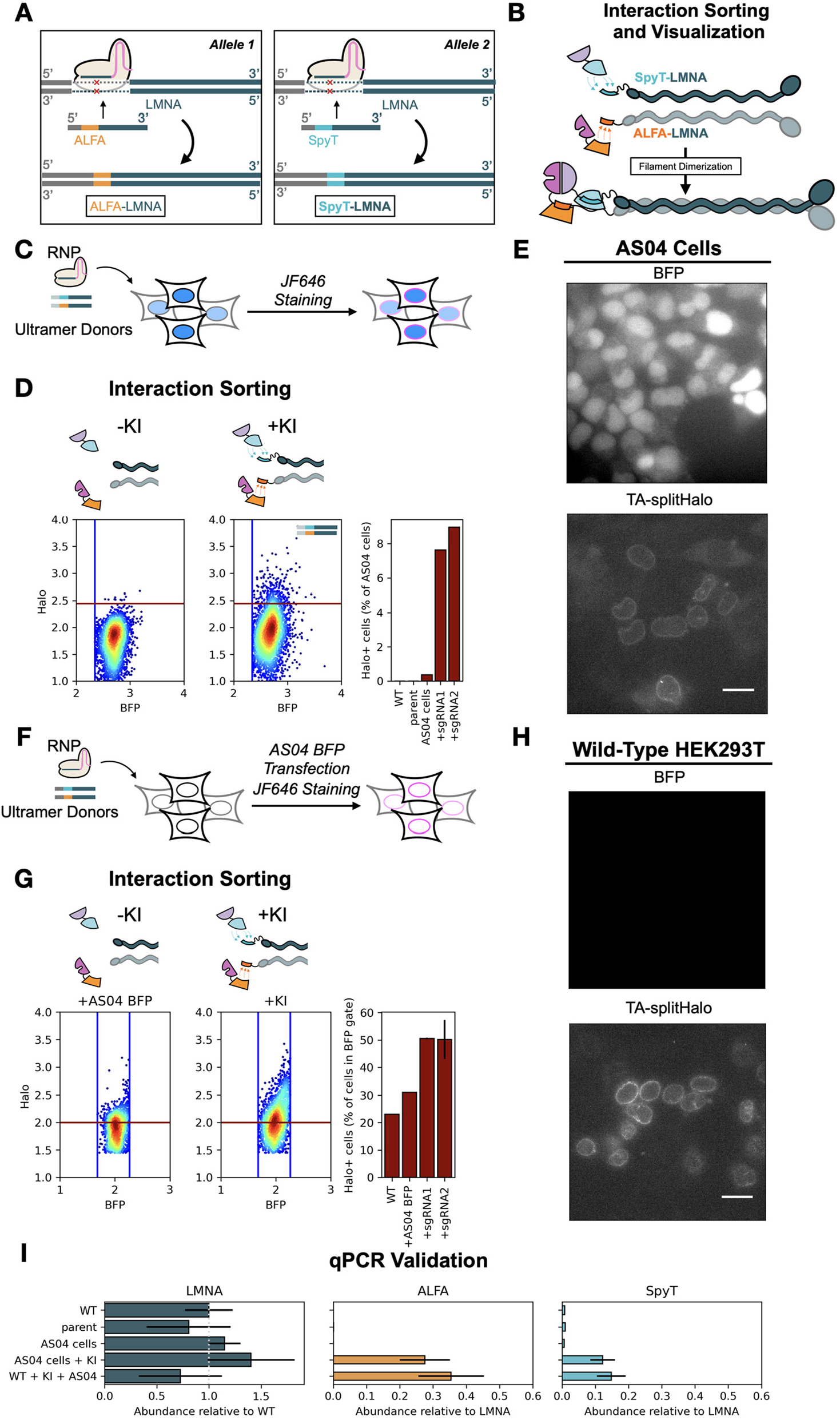
TA-splitHalo detects interactions between ALFA-LMNA and SpyT-LMNA. (A) Overview of KI strategy. (B) Schematic of lamin dimerization driving TA-splitHalo complementation. (C) Experimental workflow for ALFA/Spy TA-splitHalo interaction KIs in the AS04 cell line. (D) Flow cytometry data from KIs AS04 cells. Events (3000 per panel) are shown on log10 scale, and are gated to show mRuby-, TagBFP+ cells. The right panel graph shows the percentage of Halo+ cells for WT HEK293Ts, the parent landing pad cell line, AS04 cells without and with the KI and KI cell lines from two different sgRNAs. (E). Representative widefield microscopy images of cells sorted in (D). Scale bars: 25 μm. (F) Experimental workflow for ALFA/Spy TA-splitHalo interaction KIs in wild-type HEK293Ts. (G) Flow cytometry of KIs performed in the wild-type cells, with the AS04 BFP transfection and JF646 staining. Events (3000 per panel) are shown on log10 scale. The right panel shows the percentage of TA-splitHalo+ cells in the BFP gate. Standard deviations are shown for KIs performed in duplicates. (H). Representative widefield microscopy images of cells sorted in (G). Scale bars: 25 μm. (I) qPCR data validation of ALFA-LMNA and SpyT-LMNA KIs, with internal primers for LMNA gene and tag-specific primers for the KI alleles, showing mean ± standard deviation of quadruplicates.

Having demonstrated that our splitHalo system allows protein-protein interaction sorting in a dedicated cell line, we wanted to show that this could be achieved in a wild type background. For this purpose, we designed a reporter strategy to eliminate the high transfectants that are the source of background as seen in data from our architecture benchmarking (Figure 2). A compatible reporter would enable real-time screening and eliminate high transfectants on the FACS machine even when dampening expression by reducing the amount of transfected plasmid fails to account for all the background cells. To this end, we cloned splitHalo BFP plasmids for our selected architectures. In these plasmids, we added an mTagBFP2 reporter. mTagBFP2 has a spectral emission that does not overlap with that of JF646 and thus allows us to better sort true splitHalo positive cells.

When we performed the ALFA-LMNA + SpyT-LMNA sort in WT HEK293Ts, we transfected the AS04 BFP plasmid and set a gate on a range of BFP expression values where there is minimal Halo background in the no KI transfection control (Figure 4F). In the KI populations, we see a significant increase in Halo+ cells that mirrors our landing pad results (Figure 4G). Constraining the population of interest to cells that are minimally transfected emulates the landing pad cell line where there is only one copy of the TA-splitHalo transcriptional units. Again, we see that this signal is specific to lamin in widefield images of the sorted cells (Figure 4H).

To confirm that we are enriching cells with LMNA edits, we performed RT-qPCR on cDNA derived from RNA extracted from sorted KI and control populations for this and subsequent experiments. We used four primer pairs on each sample including one LMNA internal control and three to distinguish edited ALFA, SpyT, and GFP_11_ LMNA-specific edits (Figure S7). Compared to all controls without KIs including wild-type HEK293Ts, the parent landing pad cell line and AS04 cells, we confirmed that KI sorts enrich for both ALFA-LMNA and SpyT-LMNA KIs (Figure 4I).

### Allelic multiplexing with TA-splitHalo

Realizing that we can perform a simultaneous KI on multiple alleles, we sought to leverage the multiplexing capabilities of the two splitHalo systems for novel applications in KI enrichment. We aimed to sort cells which are GFP/Spy and ALFA/Spy TA-splitHalo compatible on the same target gene. Currently, isolating biallelic KI populations while retaining identical functionality on both loci is difficult to do without extensive clonal verification. In our special case, the dependence of the GFP/Spy system on split GFP_1-10/11_ allows us to sort the GFP_11_-SpyT KI using a traditional split GFP_1-10/11_ workflow. Thus, when we KI both GFP_11_-SpyT and ALFA-SpyT to the same gene in the same cells (Figure 5A), we can sort for each edit in a different color channel (GFP and splitHalo+ JF646 respectively), yielding cells in which TA-splitHalo can be recruited to proteins translated off multiple alleles of the same gene (Figure 5B).

**Figure 5:**
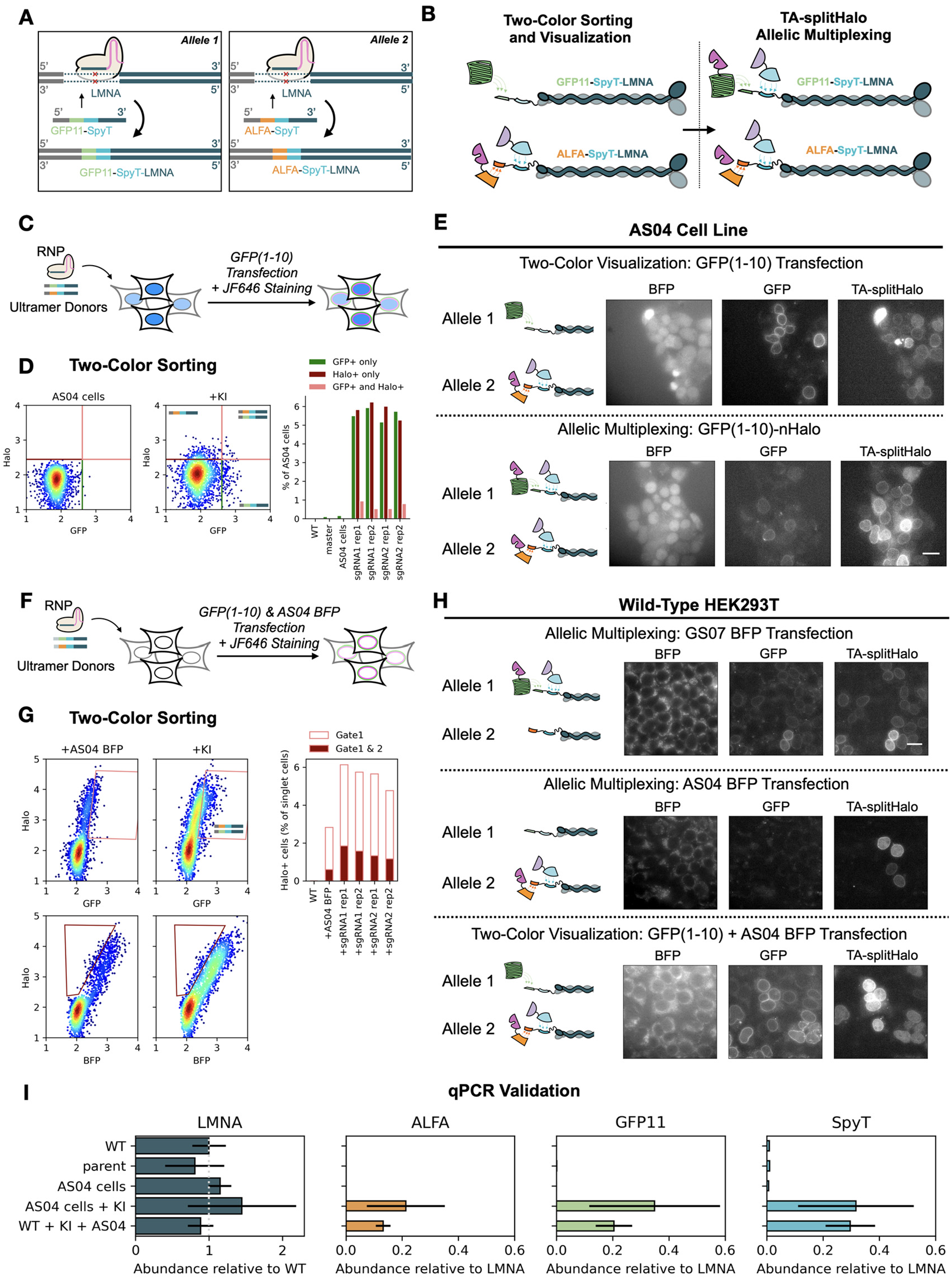
TA-splitHalo supports allelic multiplexing. (A) Overview of knock-in strategy. (B) Sorting and HaloTag functionalization of the knock-in alleles. (C) Workflow for TA-splitHalo biallelic sorting in AS04 cells. (D) Flow cytometry of KI AS04 cells with GFP_1-10_ transfection and JF646 staining. Events (2000 per panel) are shown on log10 scale, and have been gated to show mRuby-, TagBFP+ single cells. The bar graph shows the percentage of cells above thresholds for GFP, TA-splitHalo, or both in WT HEK293T, the parent landing pad cell line and AS04 cells without and with the KI. KIs were repeated in two separate wells (rep1 and rep2) with two gRNAs (sgRNA1 and sgRNA2) each targeting the N-terminal of LMNA. (E) Representative widefield microscopy images of cells sorted in (D) containing both edits. Scale bars: 25 μm. (F) Workflow for TA-splitHalo biallelic sorting in the WT HEK293T cells. (G) Flow cytometry of WT or KI HEK293T transiently co-transfected with AS04 BFP plasmid and GFP_1-10_ plasmid, followed by JF646 staining. Events (2000 per panel) are shown on log10 scale and are gated on show singlet cells. The left panel is a control showing unedited cells. The center panel shows wild-type cells after the KI, GFP_1-10_ and AS04-BFP cotransfection, and JF646 staining. Cells inside polygon gate were sorted to enrich for GFP+ AND Halo+ cells. The bar graph shows the percentage of Halo+ cells for WT HEK293Ts without and with the cotransfection compared to cells after the KI and cotransfection. Experiments were repeated in two separate wells (rep1 and rep2) for two RNAs (sgRNA1 and sgRNA2) each targeting the N-terminal of LMNA. (H) Representative widefield images of cells sorted in (G). Scale bars: 25 μm. (I) qPCR validation of ALFA-SpyT-LMNA and GFP11-SpyT-LMNA KIs, with internal primers for LMNA gene and tag-specific primers for the KI alleles, showing mean ± standard deviation of quadruplicates.

Like with the protein-protein interaction sorts, we first performed this sort using the AS04 landing pad. This cell line already contains the detection components for the ALFA/Spy TA-splitHalo system at optimal concentrations and cells were gated for the presence of BFP and absence of mRuby as above to ensure integration of the splitHalo detection fragments,so GFP_1-10_ was the only transfection needed for the two-color biallelic sort. Performing a LMNA KI with a mixture of two ultramer donors, one containing GFP_11_-SpyT and the other containing ALFA-SpyT, we expect cells that are GFP+ and Halo+ to have both KIs on separate alleles of the LMNA gene (Figure 5C). As well as GFP+ or Halo+ cells, we see an enrichment of Halo+ GFP+ cells in the KI population (Figure 5D). After sorting this population, we demonstrated multiplexing of both splitHalo systems by transfecting GFP_1-10_-nHalo. The addition of this component recapitulates the GS02 architecture in a cell line that already contains AS04, because the SpyC-cHalo fragment is shared by both of architectures (Figure S1). In cells which take up the plasmid, we expect to see nuclear envelope signal in both colors corresponding to the two edited alleles and an increase in splitHalo signal due to the presence of both TA-splitHalo systems. Our widefield images confirm the expected split GFP_1-10/11_ and TA-splitHalo lamin signal and show a clear increase in signal in transfected cells (Figure 5E). As with the protein-protein interaction experiments, we grew the enriched cells from each population. We generated cDNA from these samples and analyzed the relative fraction of edits in each population by qPCR. These results confirm enrichment of all three tags in these cells compared to pertinent controls (Figure 5F).

We also performed similar KIs on LMNA in wild-type HEK293Ts. Before sorting we transfected the cells (+/-KI) with GFP_1-10_ and AS04 BFP. GFP_1-10_ was used to sort GFP+ cells containing the GFP_11_-SpyTag KI while AS04 BFP is used to sort Halo+ cells containing the ALFA-SpyT KI (Figure 5G). To sort the population with both edits, we used a “true-splitHalo positive” gate to account for the proportional increases of TA-splitHalo signal in high transfectants and a nested gate to sort for GFP+ cells (Figure 5G). Widefield imaging shows that we can use GFP/Spy and ALFA/Spy TA-splitHalo systems in this population as well as visualize protein from both alleles using the same transfection we used to sort (Figure 5H). Again, qPCR validated enrichment of ALFA, GFP_11_, and SpyT (Figure 5I).

### TA-splitHalo Exemplifies and Enables TASEC Approaches

We introduce TASEC, a technique that employs short peptide tags to recruit split enzymes, enabling complex interfacing with target proteins with minimal scarring. Specifically, we illustrate how to engineer a TASEC system and leverage its strengths in CRISPR/Cas9-mediated KIs. The utilization of TASEC enables us to reconstruct enzymes with desired functions on any endogenous target conditional upon a specific genetic edit. Here, we applied this strategy to develop TA-splitHalo.

TA-splitHalo proved to be an ideal platform to demonstrate the strengths of the TASEC approach. It is a scalable platform that expands our capabilities for enriching KI cells and generates versatile cell lines that can exploit the full suite of HaloTag applications. Additionally, TA-splitHalo offers a rapid, non-destructive method to select and validate tandem tagged KI cells. These cell lines could then be used for architecture tests of any TASEC system. For example, *Renilla Luciferase* has ~35% homology to the HaloTag and its split may be interchangeable with splitHalo once a successful TA-splitHalo system has been identified ^6^. Other existing split enzymes that could be tested as TASEC systems in tandem tagged cell lines include split-TEV protease ^16^, split-Cre recombinase ^17^, split-Firefly luciferase ^18^, split-DamID ^19^ and split-esterase ^20^. Though we have optimized the system in human cell lines, the TA-splitHalo systems we describe should be applicable in model systems across all three kingdoms (eukaryotes, prokaryotes, and archaea).

The flow cytometry-based approach we used to decipher working TASEC architectures is applicable to any split enzyme with a fluorescence readout. From this approach, we derived two different TA-splitHalo systems from our architecture scanning that yield unique benefits. The GFP/Spy TA-splitHalo system incorporates a split-FP as one of the tag/binder pairs. This can be used to increase stringency while sorting and retain the convenience of fluorescence imaging of endogenous targets when using non-fluorescent HaloTag ligands. The ALFA/Spy TA-splitHalo system provides a way to recruit splitHalo with no extraneous fluorophores. The system yields “turn-on” Halo-tag fluorescence where cells remain dark in all channels even after full complementation of the ALFA/Spy architecture. ALFA Tag and NbALFA mutants are also excellent templates for developing orthogonal mutants and further multiplexing capabilities without the use of splitFPs that restrict applications in specific color channels.

### TA-splitHalo Expands Utility of CRISPR/Cas Knock-In Methods

Our demonstration of detecting a protein-protein interaction using TA-splitHalo provides an example of how TASEC systems can be used to study relationships between endogenous molecules. For investigating characterized interaction partners, TA-splitHalo provides a way to translate these studies into environments with high autofluorescence like organoids, embryos, and animal models due to the possibility to use long wavelength dyes^21^. By varying the concentration of Halo dye, TA-splitHalo could be used to study protein-protein interactions at the single molecule level, with limiting dye, or at the macro level, with saturating dye. When screening for unknown interaction partners, TA-splitHalo can be used in an unbiased screen to sort, validate, and possibly purify interaction partners. In the future, we can look to place a pair of TASEC tags on adaptor proteins that bind to specific DNA and RNA sequences like noncutting variants of Cas9^22^ and Cas13^23^ respectively. In this way, we can generate TASEC functionality driven by the presence of specific DNA or RNA sequences.

We have also shown that TA-splitHalo enables the sorting of complex populations by isolating biallelic KIs using multiplexing of the two TA-splitHalo systems. Employing both splitHalo systems simultaneously in a single round of FACS, we have bypassed the clonal selection and multiple genotyping steps that traditionally made this process laborious. Furthermore, if we use TA-splitHalo tagging schemes solely for enrichment, other functional sequences of interest can be added to each donor strand. Resulting cell lines therefore would contain either the same KIs on multiple alleles or varied KIs on each allele. This is a particularly important advance when tagging both alleles of a gene with a protein or peptide tag that is not detectable via FACS sorting. Additionally, the ability to sort cells with KIs on both alleles allows for manipulation of each allele separately or together in the same cell line through RNAi or protein fusions containing the TA-splitHalo binders. This is important for applications where there is a difference between perturbing one allele or both alleles. Ability to sort biallelic KIs in this fashion also empowers studying patient-derived cellular models of genetic disease where one allele is altered and behaves differently than the other. Finally, methods to separate genetically modified cells by number of alleles edited will be an important quality control for cell therapies in the future ^24^.

While TA-splitHalo is a notable advance for high-throughput sorts of complicated KI populations, a key feature is that the library of compatible ligands maximizes the potential of the sorted cell lines. Since the splitHalo ligand and saturation level can be decided on after a KI occurs and just prior to any application, the most appropriate ligand can be strategically selected each time the TA-splitHalo system is employed in the same cell line. For example, in protein labelling applications, TA-splitHalo is the first platform that outperforms the background adjusted brightness of GFP while also retaining the cost-effective workflows of split FPs when using JF646 (Figure S8). This property should allow a wider range of the human proteome to be sorted and imaged. Halo dyes in other channels can be selected to work around other fluorophores. This attribute would be valuable for flexibility in multicolor flow cytometry panels and imaging experiments. Finally, the library of available HaloTag ligands also includes molecules to facilitate purification^25^ and degradation^26,27^ of target proteins that widens the range of experiments possible with TA-splitHalo.

In conclusion, TA-splitHalo provides a modular, minimalist, scalable means to sort traditional or complex KI populations with a growing library of HaloTag ligands, making the system highly versatile. It also provides a blueprint for applying a TASEC approach to CRISPR/Cas9-mediated KIs and a path to developing new TASEC systems that can generate custom readouts linked to expression of native macromolecules or interactions between them with short peptide tags.

## Methods

### Cloning

We generated ‘part’ vectors, ‘expression’ vectors, and ‘landing pad’ vectors following the Mammalian Toolkit (MTK) approach ^28^.

The 10 μL reactions to generate part vectors consisted of 40 fmol insert DNA clean of BsaI and BsmBI restriction sites, 20 fmol MTK part vector backbone, 10x T4 Ligase Buffer (NEB B0202S), Esp31 (NEB R0734S/L), and T7 DNA Ligase (M0318S/L). The reactions were cycled between digestion at 37 ºC for 2 minutes and ligation at 25 ºC for 5 minutes. From the resulting reaction mixture, 1 μL was transformed into MachI *E. coli* (QB3 Macrolab) and colonies lacking GFP expression were selected for amplification and sequencing verification.

To streamline cloning of the expression vectors, transcriptional unit specific CDS backbones were generated by adding the requisite connector sequences, a PGK promoter, a BGH terminator and poly(A) to the original MTK assembly backbone, also known as pYTK095 (Addgene #65202). With these backbones, we improved workflows by reducing the number of inserts needed to generate new assemblies. Expression vectors were generated in 10 μL reactions containing 20 fmol CDS backbone, 40 fmol of each part insert, 10x T4 Ligase Buffer, BsaI-HF v2.0 (NEB R3733S/L), and T7 DNA Ligase with the same cycling conditions as the part vectors.

Landing pad (LP) vectors were generated similarly to part vectors in 10 μL reactions with 20 fmol MTK landing pad entry backbone (Addgene #123932), 40 fmol of each expression vector plasmids, 10x T4 Ligase Buffer (NEB B0202S), Esp31 (NEB R0734S/L), and T7 DNA Ligase (M0318S/L) with the same cycling conditions as the part vectors. For generating landing pad vectors from expression vectors without the correct overhangs, an oligonucleotide stuffer was used to complete the overhangs.

TA-splitHalo BFP plasmids were made in 10 μL reactions comprising 20 fmol Kanamycin ColE1 digested backbone, 40 fmol TA-splitHalo fusion expression vectors, 40 fmol PGK-mTagBFP2 expression vector, 10x T4 Ligase Buffer (NEB B0202S), Esp31 (NEB R0734S/L), and T7 DNA Ligase (NEB M0318S/L) with the same cycling conditions as the part vectors.

### Transfection of HeLa cells in 8-well chamber flasks for TA-splitHalo Architecture Benchmarking

For Figure 1 transfections, HeLa cells were seeded in an 8-well chamber flask at 20k cells per well in 225 μL DMEM +1% Penicillin/Streptomycin (P/S) 10% Fetal Bovine Serum (FBS). 160 ng of each TA-splitHalo architecture plasmid was cotransfected with 80 ng of mCherry bait plasmid with 0.7 μL FuGENE HD. This corresponded to a 1:1 molar ratio. After an overnight incubation, samples were stained with 10 nM JF646 in 100 μL Phenol Red-free DMEM +1% Penicillin/Streptomycin +10% Fetal Bovine Serum. Flow cytometry was performed the day after overnight staining.

### Seeding and Transfection of HEK293T KIs for TA-splitHalo Sorting

In all experiments 6-well chamber flasks were seeded with 300k pre-sorted KI cells and controls were seeded in 2 mL of DMEM +1% P/S 10% FBS.

For Figure 4G, 180 fmol of AS04 BFP plasmid was transfected with 2.8 μL FuGENE HD in each well containing control HEK2993Ts and pre-sorted LMNA KI cells.

For Figure 5D in AS04 cells, 600 fmol of GFP1-10 plasmid was transfected with 9.3 μL FuGENE HD in each well containing control AS04 cells and pre-sorted LMNA KI AS04 cells.

For Figure 4G, 600 fmol of GFP1-10 and 180fmol of AS04 BFP plasmid were cotransfected with 9.3 μL FuGENE HD in each well containing control AS04 cells and pre-sorted LMNA KI AS04 cells.

In all cases, cells were stained in 10nM JF646 in 1mL of Phenol Red-free DMEM +1% P/S 10% FBS after an overnight transfection. Cells were FACS sorted the day after staining.

### Seeding and Transfecting HEK293Ts for TA-splitHalo Imaging

In all experiments, 8-well chamber flasks were pre-treated with poly-L-lysine seeding. For AS04 BFP imaging in Figure 4H, 20k HEK293T cells containing ALFA-LMNA SpyT-LMNA KIs were seeded in each well. After incubation overnight, 15 fmol AS04-BFP was transfected with 0.7 μL FuGENE HD.

For AS04 cell imaging in Figure 5E, 20k sorted AS04 cells containing ALFA-LMNA SpyT-LMNA KIs were seeded in each well. After incubation overnight, we performed 50fmol transfections of GFP1-10 and GFP1-10-nHalo with 0.7 μL FuGENE HD in different wells.

For AS04 cell imaging in Figure 5E, 20k sorted HEK293T cells containing ALFA-LMNA SpyT-LMNA KIs were seeded in each well. After incubation overnight, we performed 15 fmol GS07 BFP, 15 fmol AS04 BFP, and 50fmol GFP1-10 + 15 fmol AS04 BFP transfections with 0.7 μL FuGENE HD in different wells.

After each of these transfections, Cells were stained with 10 nM JF646 after an overnight incubation in 100 μL Phenol Red-free DMEM +1% Penicillin/Streptomycin 10% Fetal Bovine Serum and imaged the subsequent day.

### Lamin A/C gRNA IVT Template Synthesis

The IVT template for LMNA gRNA was made by PCR. The reactions are done in a 100 μL reaction containing 50 μL 2x Phusion MM (ThermoFischer F531L), 2 μL ML557+558 mix at 50 μM, 0.5 μL ML611 at 4 μM, 0.5 μL of each gene-specific oligo at 4 μM, and 47 μL DEPC H_2_O. The PCR product was purified using a Zymo DNA Clean and Concentrator Kit (Zymo Research D4014). Sequences for these primers and thermocycling conditions are given in Figure S7)

### Lamin A/C gRNA Synthesis

IVT was carried out using the HiScribe T7 Quick High Yield RNA Synthesis Kit (NEB E2050S) with the addition of RNAsin (Promega N2111). Purification of mRNA was performed using the RNA Clean and Concentrator Kit (Zymo Research R1017). gRNA was stored at −80°C immediately after measuring concentration and diluting to 130 μM.

### Generation of Split-Halo Landing Pad Detection Cell Lines

The split-GFP, and split-Halo Landing Pad HEK293Ts, were generated from a published landing pad parent cell line^28^ seeded at 100k cells in a 12-well plate. To each well, 600 ng of BxbI Integrase Expression Vector (Addgene #51271) and 600 ng of each landing pad donor plasmid were co-transfected. Once cells are confluent, cells were split once and seeded in a T25 flask, and blasticidin (Gemini Bio-Products 400-165P) was added at 5 μg/mL for selection prior to FACS sorting integrated cell lines.

### Cas9 HDR Knock-Ins

The day prior to performing the KI, 2.5 million HEK293Ts were treated with 200 ng/mL nocodazole and seeded at 250k cells/mL in 10 mL DMEM media (Sigma-Aldrich M1404) before incubation overnight for 15-18h prior to nucleofection.

The next day, RNPs were generated in 10 μL reactions consisting of 1 μL sgRNA at 130 μM, 2.5 μL purified Cas9 at 40 μM, 1.5 μL HDR template at 100 μM, 2 μL 5x Cas9 Buffer, and DEPC H_2_O up to 10 μL. HDR template ultramer sequences synthesized from IDT are given in Table S1.

In a sterile PCR or microcentrifuge tube, Cas9 Buffer, DEPC H_2_O, and sgRNA were mixed and incubated at 70°C for 5 min to refold the gRNA. During this step, 10 μL aliquots of purified Cas9 at 40 μM was thawed on ice. Next, 2.5 μL Cas9 protein was slowly added to the diluted sgRNA in Cas9 buffer and incubated at 37°C for 10 min. Finally,1.5 μL of each ultramer donor was to the RNP mix and all samples were kept on ice until ready for nucleofection.

For efficient recovery post-KI, a 24-well plate with 1 mL media per well was incubated in a 37°C. An appropriate amount of supplemented Amaxa solution corresponding to the number of KIs to be performed was prepared room temp in the cell culture hood. For each sample 16.4 μL SF solution and 3.6 μL supplement was added to an Eppendorf tube for a total of 20 μL per KI. Amaxa nucleofector instruments/computers were then turned on and kept ready for nucleofection.

Nocodazole-treated cells were harvested into a sterile Falcon tube and counted. A volume equivalent to 200k cells per KI was transferred to another Falcon tube and centrifuged at 500g for 3 min. Remove supernatant containing nocodazole-treated media and resuspend in 1 mL PBS to wash. The cells were centrifuged again at 500g for 3 min. PCR tubes containing RNPs were brought into TC hood.

Cells were resuspended in supplemented Amaxa solution at a density of 10k cells/μL. 20 μL of the cell resuspension was added to each 10 μL RNP tube. The cell/RNP mix was pipetted into the bottom of the nucleofection plate. The nucleofection was carried out on a Lonza 96-Well shuttle Device (Lonza AAM-1001S) attached to Lonza 4D Nucleofector Core Unit (Lonza AAF-1002B). Cells were nucleofected using CM-130 program and recovered using 100 μL media from the pre-warmed 24-well plate and transferred to the corresponding well.

Once cells reached 80% confluence in the smaller vessel, they were transferred first to a 6-well plate and then to a T25 flask. Cells were FACS sorted after a week of maintaining and expanding the pre-sorted KI population to reach optimal cell numbers and Cas9-mediated cutting and repair.

### Cell Line Genotyping

Genomic DNA was prepared from 1 million cells using the Monarch Genomic DNA Purification Kit (NEB, #T3010G). Diagnostic PCR was then carried out followed by gel extraction (NucleoSpin) and Sanger Sequencing (Quintara Biosciences).

### Confocal imaging

Cells were imaged on a Nikon Ti Microscope equipped with a Yokagawa CSU22 spinning disk confocal and an automated Piezo stage. We used a CO_2_- and temperature-controlled incubator it is ideal for live specimen imaging. Our laser lines were 405nm, 491nm, 561nm, 640nm. Pixel binning was set at 2×2.

### Widefield imaging

All widefield imaging was performed on a Nikon Ti-E microscope equipped with a motorized stage, a Hamamatsu ORCA Flash 4.0 camera, an LED light source (Excelitas X-Cite XLED1), and a 60X CFI Plan Apo IR water immersion objective. All downstream image analysis was performed in ImageJ.

### qPCR

Total RNA was extracted from 1 million cells using the Monarch Total RNA Miniprep Kit (NEB, #T2010S). We prepared cDNA from 1 μg of extracted RNA using LunaScript^®^ RT SuperMix Kit (NEB, #E3010). No Template and No Reverse Transcriptase controls (NTC and NRT) were performed in parallel to cDNA preparations. We set up qPCR plates using 0.5 μl of each 20 μl cDNA sample,10 μl 2x Maxima SYBR Green qPCR Master Mix (Thermo Scientific K0221), and optimized primer pairs corresponding to SpyT-specific, GFP_11_-specific, and ALFA-specific LMNA KIs. We also ran a primer set specific to the wild-type LMNA gene for a positive control and reference marker.

For standard curves, we cloned plasmids containing sequences corresponding to all edited and unedited versions of the LMNA gene. RT-qPCR was performed on QuantStudio™ 5 Real-Time PCR System. These primer sequences and a schematic for our qPCR experiments are shown in Figure S7.

### Flow cytometry analysis and cell sorting

FACS sorting and flow cytometry was performed on a BD FACSAria II in the Laboratory for Cell Analysis at UCSF. BFP or mTagBFP2 signal was measured using the 405 nm laser with a 450/50 bandpass filter, GFP signal was measured with the 488 nm laser and 530/30 bandpass filter, mRuby or mCherry signal was measured using the 561 nm laser and 610/20 bandpass filter and TA-splitHalo signal using JF646 was measured with the 633 nm laser with a 710/50 bandpass filter. Files in the .fcs format were exported from the BD FACS Aria II were analyzed in Python using our altFACS package (https://pypi.org/project/altFACS/).

## Supporting information

Supplementary Data (Nucleic Acid Sequences)

## Data Availability

Raw data and Jupyter notebooks used to prepare this manuscript are available on our GitHub page (https://github.com/BoHuangLab).

## Acknowledgements

We thank Zene Matsuda for the original splitHalo plasmids and Luke Lavis for the JF646 HaloLigand. Thanks to Alejandro Ramirez-Apodaca and Aivy David for excellent lab management, Sarah Elmes for help with FACS, DeLaine Larsen and Kari Herrington and the Nikon Imaging Center for help with confocal imaging, Alain Bonny and the El-Samad lab for help with the MTK system and landing pad parent cell line. Thanks to Jonathan Weissman for inspirational discussions and to Nairi Hartooni, Taia Wu and Shengyan Jin for their help early in this project.

## References

(1) Cabantous, S.; Terwilliger, T. C.; Waldo, G. S. Protein Tagging and Detection with Engineered Self-Assembling Fragments of Green Fluorescent Protein. Nature Biotechnology 2005, 23 (1), 102–107. https://doi.org/10.1038/nbt1044.

(2) Kamiyama, D.; Sekine, S.; Barsi-Rhyne, B.; Hu, J.; Chen, B.; Gilbert, L. A.; Ishikawa, H.; Leonetti, M. D.; Marshall, W. F.; Weissman, J. S.; Huang, B. Versatile Protein Tagging in Cells with Split Fluorescent Protein. Nature Communications 2016, 7 (1), 11046. https://doi.org/10.1038/ncomms11046.

(3) Feng, S.; Sekine, S.; Pessino, V.; Li, H.; Leonetti, M. D.; Huang, B. Improved Split Fluorescent Proteins for Endogenous Protein Labeling. Nature Communications 2017, 8 (1), 370. https://doi.org/10.1038/s41467-017-00494-8.

(4) Feng, S.; Varshney, A.; Coto Villa, D.; Modavi, C.; Kohler, J.; Farah, F.; Zhou, S.; Ali, N.; Müller, J. D.; Van Hoven, M. K.; Huang, B. Bright Split Red Fluorescent Proteins for the Visualization of Endogenous Proteins and Synapses. Communications Biology 2019, 2 (1), 1–12. https://doi.org/10.1038/s42003-019-0589-x.

(5) Leonetti, M. D.; Sekine, S.; Kamiyama, D.; Weissman, J. S.; Huang, B. A Scalable Strategy for High-Throughput GFP Tagging of Endogenous Human Proteins. PNAS 2016, 113 (25), E3501–E3508. https://doi.org/10.1073/pnas.1606731113.

(6) Paulmurugan, R.; Gambhir, S. S. Monitoring Protein–Protein Interactions Using Split Synthetic Renilla Luciferase Protein-Fragment-Assisted Complementation. Anal Chem 2003, 75 (7), 1584–1589.

(7) Gao, X. J.; Chong, L. S.; Kim, M. S.; Elowitz, M. B. Programmable Protein Circuits in Living Cells. Science 2018, 361 (6408), 1252–1258. https://doi.org/10.1126/science.aat5062.

(8) Ishikawa, H.; Meng, F.; Kondo, N.; Iwamoto, A.; Matsuda, Z. Generation of a Dual-Functional Split-Reporter Protein for Monitoring Membrane Fusion Using Self-Associating Split GFP. Protein Engineering Design and Selection 2012, 25 (12), 813–820. https://doi.org/10.1093/protein/gzs051.

(9) Grimm, J. B.; Brown, T. A.; English, B. P.; Lionnet, T.; Lavis, L. D. Synthesis of Janelia Fluor HaloTag and SNAP-Tag Ligands and Their Use in Cellular Imaging Experiments. In Super-Resolution Microscopy; Erfle, H., Ed.; Methods in Molecular Biology; Springer New York: New York, NY, 2017; Vol. 1663, pp 179–188. https://doi.org/10.1007/978-1-4939-7265-4_15.

(10) Grimm, J. B.; English, B. P.; Choi, H.; Muthusamy, A. K.; Mehl, B. P.; Dong, P.; Brown, T. A.; Lippincott-Schwartz, J.; Liu, Z.; Lionnet, T.; Lavis, L. D. Bright Photoactivatable Fluorophores for Single-Molecule Imaging. Nat Methods 2016, 13 (12), 985–988. https://doi.org/10.1038/nmeth.4034.

(11) Zheng, Q.; Ayala, A. X.; Chung, I.; Weigel, A. V.; Ranjan, A.; Falco, N.; Grimm, J. B.; Tkachuk, A. N.; Wu, C.; Lippincott-Schwartz, J.; Singer, R. H.; Lavis, L. D. Rational Design of Fluorogenic and Spontaneously Blinking Labels for Super-Resolution Imaging. ACS Cent Sci 2019, 5 (9), 1602–1613. https://doi.org/10.1021/acscentsci.9b00676.

(12) Shi, X.; Li, Q.; Dai, Z.; Tran, A. A.; Feng, S.; Ramirez, A. D.; Lin, Z.; Wang, X.; Chow, T. T.; Seiple, I. B.; Huang, B. Label-Retention Expansion Microscopy. BioRxiv 2019. https://doi.org/10.1101/687954.

(13) Götzke, H.; Kilisch, M.; Martínez-Carranza, M.; Sograte-Idrissi, S.; Rajavel, A.; Schlichthaerle, T.; Engels, N.; Jungmann, R.; Stenmark, P.; Opazo, F.; Frey, S. The ALFA-Tag Is a Highly Versatile Tool for Nanobody-Based Bioscience Applications. Nat Commun 2019, 10 (1), 4403. https://doi.org/10.1038/s41467-019-12301-7.

(14) Dittmer, T. A.; Misteli, T. The Lamin Protein Family. Genome Biol 2011, 12 (5), 222. https://doi.org/10.1186/gb-2011-12-5-222.

(15) Ahn, J.; Jo, I.; Kang, S.; Hong, S.; Kim, S.; Jeong, S.; Kim, Y.-H.; Park, B.-J.; Ha, N.-C. Structural Basis for Lamin Assembly at the Molecular Level. Nature Communications 2019, 10 (1), 3757. https://doi.org/10.1038/s41467-019-11684-x.

(16) Wehr, M. C.; Laage, R.; Bolz, U.; Fischer, T. M.; Grünewald, S.; Scheek, S.; Bach, A.; Nave, K.-A.; Rossner, M. J. Monitoring Regulated Protein-Protein Interactions Using Split TEV. Nat Methods 2006, 3 (12), 985–993. https://doi.org/10.1038/nmeth967.

(17) Hirrlinger, J.; Scheller, A.; Hirrlinger, P. G.; Kellert, B.; Tang, W.; Wehr, M. C.; Goebbels, S.; Reichenbach, A.; Sprengel, R.; Rossner, M. J.; Kirchhoff, F. Split-Cre Complementation Indicates Coincident Activity of Different Genes in Vivo. PLoS ONE 2009, 4 (1), e4286. https://doi.org/10.1371/journal.pone.0004286.

(18) Paulmurugan, R.; Umezawa, Y.; Gambhir, S. S. Noninvasive Imaging of Protein–Protein Interactions in Living Subjects by Using Reporter Protein Complementation and Reconstitution Strategies. Proc Natl Acad Sci U S A 2002, 99 (24), 15608–15613. https://doi.org/10.1073/pnas.242594299.

(19) Hass, M. R.; Liow, H.-H.; Chen, X.; Sharma, A.; Inoue, Y. U.; Inoue, T.; Reeb, A.; Martens, A.; Fulbright, M.; Raju, S.; Stevens, M.; Boyle, S.; Park, J.-S.; Weirauch, M. T.; Brent, M. R.; Kopan, R. SpDamID: Marking DNA Bound by Protein Complexes Identifies Notch-Dimer Responsive Enhancers. Mol Cell 2015, 59 (4), 685–697. https://doi.org/10.1016/j.molcel.2015.07.008.

(20) Jones, K. A.; Kentala, K.; Beck, M. W.; An, W.; Lippert, A. R.; Lewis, J. C.; Dickinson, B. C. Development of a Split Esterase for Protein-Protein Interaction-Dependent Small-Molecule Activation. ACS Cent Sci 2019, 5 (11), 1768–1776. https://doi.org/10.1021/acscentsci.9b00567.

(21) Heppert, J. K.; Dickinson, D. J.; Pani, A. M.; Higgins, C. D.; Steward, A.; Ahringer, J.; Kuhn, J. R.; Goldstein, B. Comparative Assessment of Fluorescent Proteins for in Vivo Imaging in an Animal Model System. MBoC 2016, 27 (22), 3385–3394. https://doi.org/10.1091/mbc.e16-01-0063.

(22) Chen, B.; Gilbert, L. A.; Cimini, B. A.; Schnitzbauer, J.; Zhang, W.; Li, G.-W.; Park, J.; Blackburn, E. H.; Weissman, J. S.; Qi, L. S.; Huang, B. Dynamic Imaging of Genomic Loci in Living Human Cells by an Optimized CRISPR/Cas System. Cell 2013, 155 (7), 1479–1491. https://doi.org/10.1016/j.cell.2013.12.001.

(23) Abudayyeh, O. O.; Gootenberg, J. S.; Essletzbichler, P.; Han, S.; Joung, J.; Belanto, J. J.; Verdine, V.; Cox, D. B. T.; Kellner, M. J.; Regev, A.; Lander, E. S.; Voytas, D. F.; Ting, Y.; Zhang, F. RNA Targeting with CRISPR–Cas13. Nature 2017, 550 (7675), 280–284. https://doi.org/10.1038/nature24049.

(24) Roth, T. L. Editing of Endogenous Genes in Cellular Immunotherapies. Curr Hematol Malig Rep 2020, 15 (4), 235–240. https://doi.org/10.1007/s11899-020-00587-0.

(25) Méndez, J. L.; Ohana, R. F.; Hurst, R.; Murphy, N.; Benink, H.; Slater, M. R.; Brueck, C.; Daniels, D. L.; Urh, M. Highly Efficient Protein and Complex Purification from Mammalian Cells Using the HaloTag® Technology. BioTechniques 2011, 51 (4), 276–277. https://doi.org/10.2144/000113767.

(26) Tovell, H.; Testa, A.; Maniaci, C.; Zhou, H.; Prescott, A. R.; Macartney, T.; Ciulli, A.; Alessi, D. R. Rapid and Reversible Knockdown of Endogenously Tagged Endosomal Proteins via an Optimized HaloPROTAC Degrader. ACS Chem. Biol. 2019, 14 (5), 882–892. https://doi.org/10.1021/acschembio.8b01016.

(27) Simpson, L. M.; Macartney, T. J.; Nardin, A.; Fulcher, L. J.; Röth, S.; Testa, A.; Maniaci, C.; Ciulli, A.; Ganley, I. G.; Sapkota, G. P. Inducible Degradation of Target Proteins through a Tractable Affinity-Directed Protein Missile System. Cell Chemical Biology 2020, 27 (9), 1164–1180.e5. https://doi.org/10.1016/j.chembiol.2020.06.013.

(28) Fonseca, J. P.; Bonny, A. R.; Kumar, G. R.; Ng, A. H.; Town, J.; Wu, Q. C.; Aslankoohi, E.; Chen, S. Y.; Dods, G.; Harrigan, P.; Osimiri, L. C.; Kistler, A. L.; El-Samad, H. A Toolkit for Rapid Modular Construction of Biological Circuits in Mammalian Cells. ACS Synth. Biol. 2019, 8 (11), 2593–2606. https://doi.org/10.1021/acssynbio.9b00322.

