## Supplementary Figures and Tables for "Versatile labeling and detection of endogenous proteins using tag-assisted split enzyme complementation"

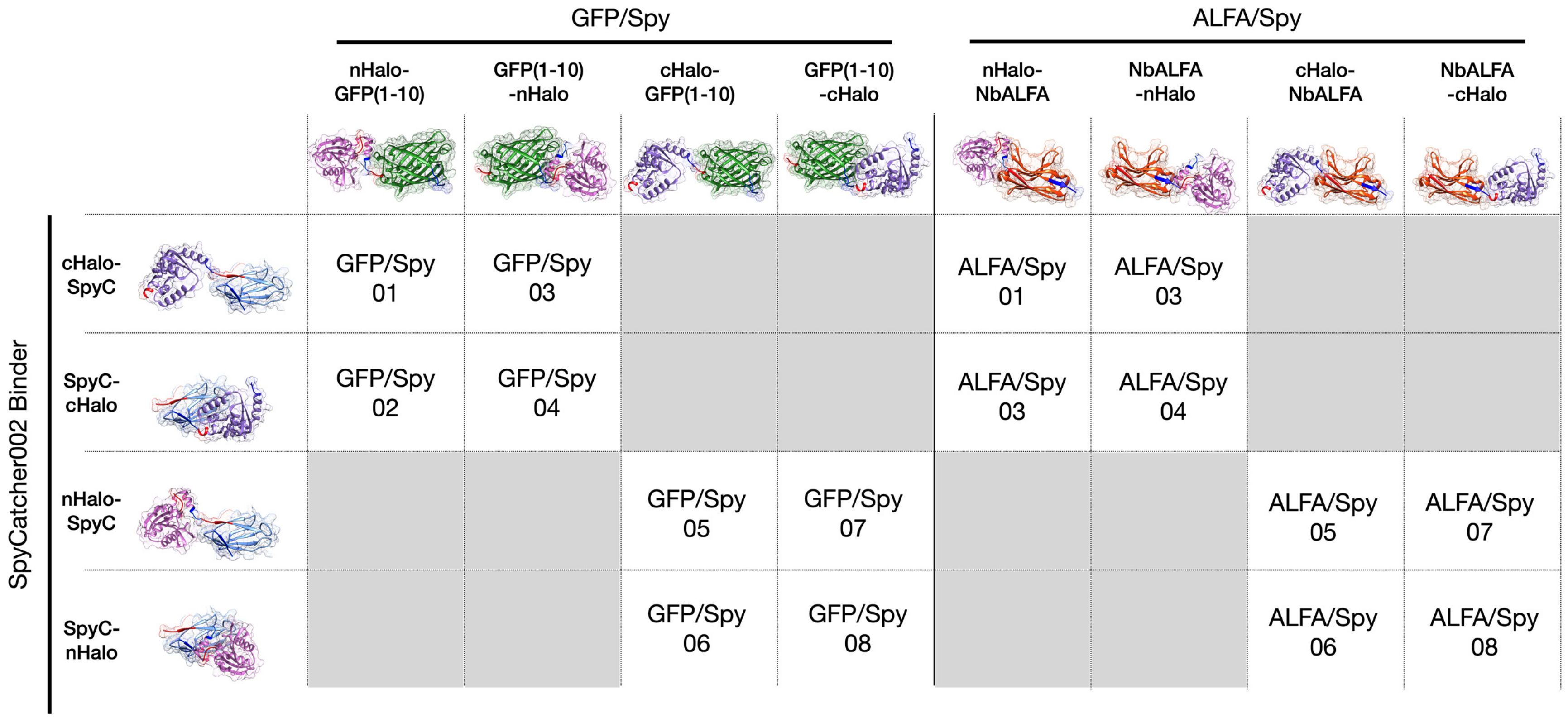


**Figure S1:** The relationship between TA-splitHalo fusions and architecture nomenclature. Numberings for each architecture define where the nHalo and cHalo components are positioned relative to each of the binders. Structural representations of TA-splitHalo fusions were made using UCSF Chimera. Components include nHalo (magenta PDB: 5UXZ), cHalo (purple PDB: 5UXZ), GFP(1-10) (green PDB 2B3P), NbALFA (PDB: orange 6I2G), and SpyCatcher002 (cyan PDB: 4MLI). N- and C-termini of all proteins are shown in red and dark blue respectively.

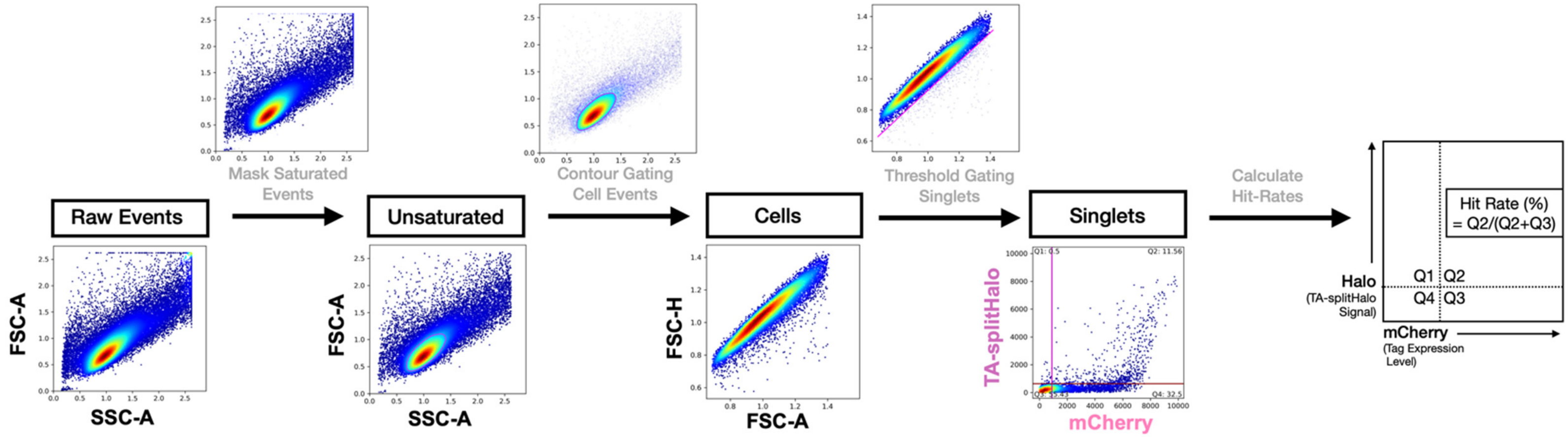


| GS01 | | | GS02 | | | GS03 | | | GS04 | | | GS05 | | | GS06 | | | GS07 | | | GS08 | | |
|---|---|---|---|---|---|---|---|---|---|---|---|---|---|---|---|---|---|---|---|---|---|---|---|
| t-value | p-value | p<.05 | t-value | p-value | p<.05 | t-value | p-value | p<.05 | t-value | p-value | p<.05 | t-value | p-value | p<.05 | t-value | p-value | p<.05 | t-value | p-value | p<.05 | t-value | p-value | p<.05 |
| -7.65 | 1.27E-02 | TRUE | -21.01 | 6.14E-04 | TRUE | -7.41 | 1.69E-02 | TRUE | -6.68 | 8.43E-03 | TRUE | -6.92 | 9.17E-03 | TRUE | -5.27 | 2.28E-02 | TRUE | -6.16 | 2.44E-02 | TRUE | -3.46 | 2.66E-02 | TRUE |
| AS01 | | | AS02 | | | AS03 | | | AS04 | | | AS05 | | | AS06 | | | AS07 | | | AS08 | | |
| t-value | p-value | p<.05 | t-value | p-value | p<.05 | t-value | p-value | p<.05 | t-value | p-value | p<.05 | t-value | p-value | p<.05 | t-value | p-value | p<.05 | t-value | p-value | p<.05 | t-value | p-value | p<.05 |
| -2.95 | 6.2E-02 | FALSE | -9.83 | 6.79E-04 | TRUE | -19.63 | 5.78E-05 | TRUE | -13.33 | 3.28E-03 | TRUE | -7.62 | 1.4E-02 | TRUE | -8.43 | 5.4E-03 | TRUE | -6.06 | 4.9E-03 | TRUE | -4.98 | 3.80E-02 | TRUE |

**Figure S2:** Workflow diagram showing flow cytometry analysis for Figure 2 with the altFACS python package. Raw events from .fcs files are processed by eliminating saturating events, gating for singlet cell events, and deriving hit rates by assessing the fraction of TA-splitHalo+ mCherry+ cells in the total mCherry+ population. The table shows the results of Welch's unequal variance t-test for the data in Figure 2.

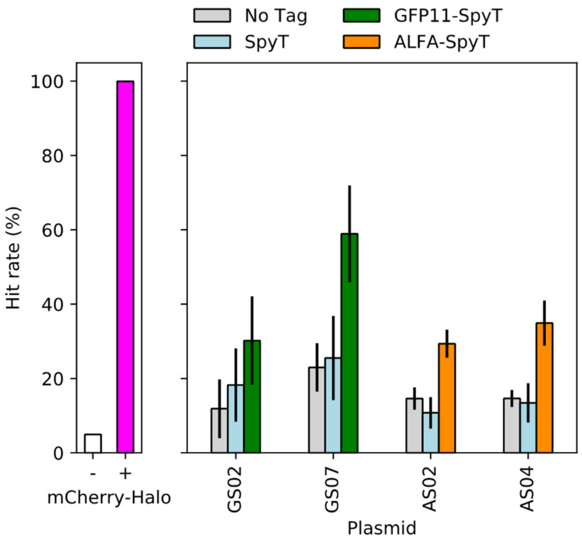


| GFP/Spy 02 | | | | | | | | |
|---|---|---|---|---|---|---|---|---|
| 0 vs. 1 Tag(s) | | | 0 vs. 2 Tags | | | 1 vs. 2 Tag(s) | | |
| t-value | p-value | p<.05 | t-value | p-value | p<.05 | t-value | p-value | p<.05 |
| -0.87 | 0.43 | FALSE | -2.22 | 0.10 | FALSE | -1.34 | 0.25 | FALSE |

| GFP/Spy 07 | | | | | | | | |
|---|---|---|---|---|---|---|---|---|
| 0 vs. 1 Tag(s) | | | 0 vs. 2 Tags | | | 1 vs. 2 Tag(s) | | |
| t-value | p-value | p<.05 | t-value | p-value | p<.05 | t-value | p-value | p<.05 |
| -0.34 | 0.76 | FALSE | -4.28 | 0.024 | TRUE | -3.35 | 0.03 | TRUE |

| ALFA/Spy 02 | | | | | | | | |
|---|---|---|---|---|---|---|---|---|
| 0 vs. 1 Tag(s) | | | 0 vs. 2 Tags | | | 1 vs. 2 Tag(s) | | |
| t-value | p-value | p<.05 | t-value | p-value | p<.05 | t-value | p-value | p<.05 |
| 1.30 | 0.27 | FALSE | -5.29 | 7.1E-03 | TRUE | -5.69 | 4.9E-03 | TRUE |

| ALFA/Spy 04 | | | | | | | | |
|---|---|---|---|---|---|---|---|---|
| 0 vs. 1 Tag(s) | | | 0 vs. 2 Tags | | | 1 vs. 2 Tag(s) | | |
| t-value | p-value | p<.05 | t-value | p-value | p<.05 | t-value | p-value | p<.05 |
| 0.36 | 0.74 | FALSE | -5.40 | 0.02 | TRUE | -4.62 | 0.01 | TRUE |

**Figure S3:** Bar plot investigating the TA-splitHalo tag-independent background. Compared to mCherry-HaloTag conditions without (-) and with (+) JF646 (left panel), selected architectures (right panel) have similar quantities of TA-splitHalo background when mCherry is untagged (grey) or tagged with only SpyT (blue). This implies that background is not driven by any single tag but rather by spontaneous complementation of TA-splitHalo fusions. For bars in the righthand panel, n=3 and errors represent standard deviation. The table shows the results of Welch's unequal variance t-test. With two tags (green and orange), we see statistically higher TA-splitHalo complementation and signal demonstrating specificity.

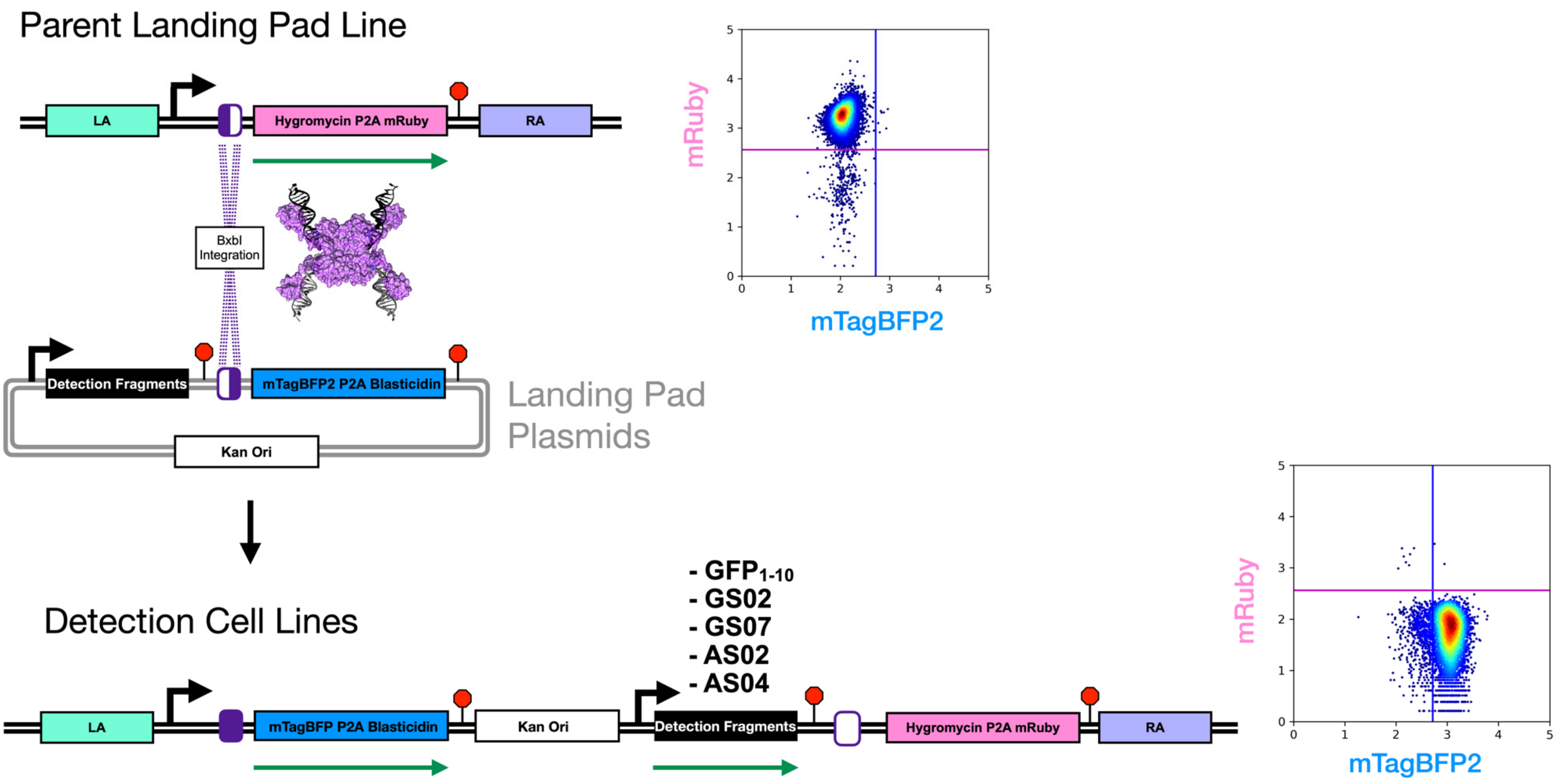


**Figure S4:** Isogenic cell lines were made from a parent cell line containing a Hygromycin-P2A-mRuby (pink) cassette at the AAVS1 safe harbor locus (top right). Flow cytometry confirms these cells as mRuby+ mTagBFP- (top right). Through BxbI integration, landing pad plasmids were integrated at this site (center). Upon successful integration, the selection/reporter cassette swaps to mTagBFP2-P2A-Blasticidin (blue) and detection component cargo will be expressed from a single genomic copy (black) (bottom left). Flow cytometry data shows an example of integrant cells that are mRuby- mTagBFP+ (bottom right). The left and right arms of the AAVS1 locus are shown in sea green and lavender. A structural representation of the serine integrase strand exchange between attB and attP sites (purple /is shown in magenta (PDB: 1ZR4). Promoters are shown with black arrows, ORFs that are transcribed and translated are shown with green arrows, and terminators are depicted as red stop signs.

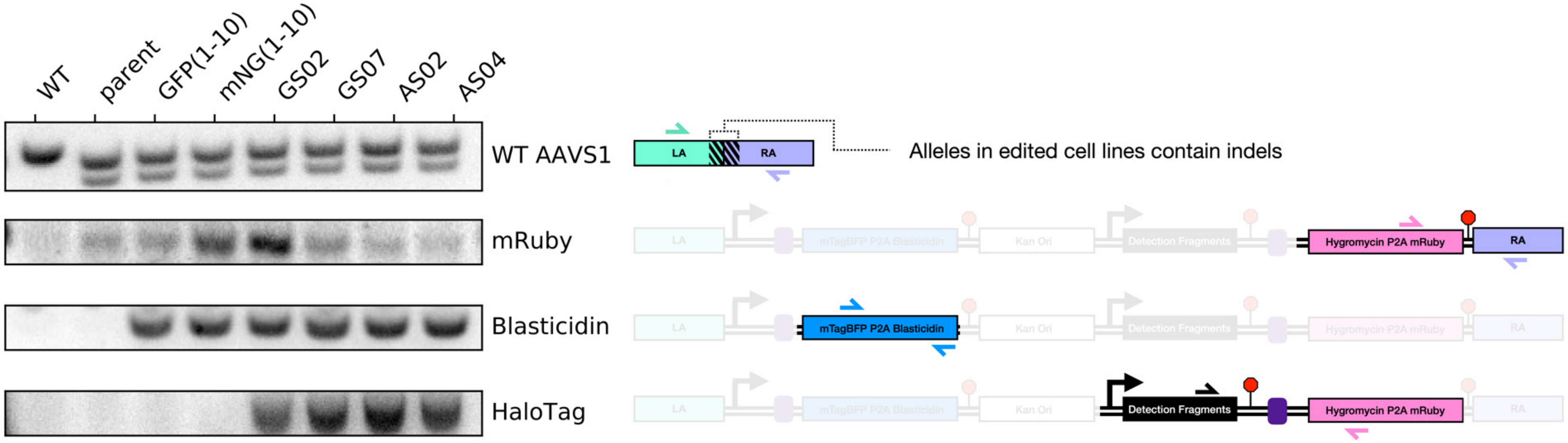


| Primer Set | Fwd Primer | Fwd Primer Sequence | Rev Primer | Rev Primer Sequence |
|---|---|---|---|---|
| **WT AAVS1** | AAVS1_Fwd | CTGGACAACCCCAAAGTACC | AAVS1_Rev | AGATGGCTCCAGGAAATGG |
| **mRuby** | mRuby_Fwd | TCTTGTGCCCAGGAGAGC | mRuby_Rev | CCCAATATCAGGAGACTAGGAAGG |
| **Blasticidin** | Blast_Fwd | CCCTCATTGAAAGAGCAACG | Blast_Rev | CCTTCACTATGGCTTTGATCC |
| **HaloTag** | Halo_Fwd | GTCGAGATGGACCATTACCG | Halo_Rev | GGGAGATGCAATAGGTCAGG |

**Figure S5:** Genotyping single integrant cell lines (left). Genomic DNA PCRs were performed to amplify the AAVS1 locus present in all cell lines (1st panel), mRuby present in each landing pad parent derived lines (2nd panel), blasticidin integrant present in each integrant line (3rd panel), and Halo-specific integrants present in TA-splitHalo detection cell lines (4th panel). The mNG(1-10) cell line made in parallel was not used in this study. Schematics defining locations of primers (black arrows) for each PCR are shown (right).

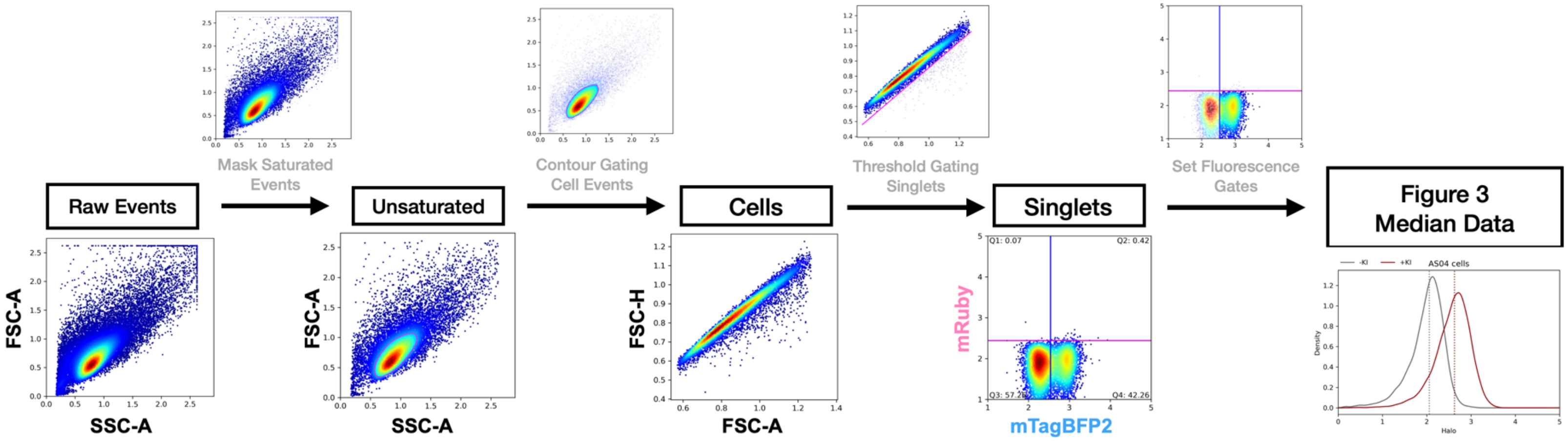


| KI Cell Line | Channel | SBR | KS value | p-value | p<.05 |
|---|---|---|---|---|---|
| $GFP_{1-10}$ | 488nm (GFP) | 1.86 | 0.58 | 0 | TRUE |
| | 646nm (TA-splitHalo) | 0.99 | 0.023 | 0.11 | FALSE |
| GS02 | 488nm (GFP) | 1.19 | 0.20 | 9.01E-51 | TRUE |
| | 646nm (TA-splitHalo) | 1.86 | 0.34 | 3.86E-149 | TRUE |
| GS07 | 488nm (GFP) | 1.26 | 0.25 | 2.9E-175 | TRUE |
| | 646nm (TA-splitHalo) | 2.05 | 0.29 | 1.39E-248 | TRUE |
| AS02 | 488nm (GFP) | 1.02 | 0.042 | 6.35E-06 | TRUE |
| | 646nm (TA-splitHalo) | 3.41 | 0.59 | 0 | TRUE |
| AS04 | 488nm (GFP) | 1.04 | 0.046 | 1.06E-07 | TRUE |
| | 646nm (TA-splitHalo) | 3.72 | 0.61 | 0 | TRUE |

**Figure S6:** Workflow diagram depicting flow cytometry analysis for Figure 3 with the altFACS python package. Raw events from .fcs files are processed by eliminating saturating events, gating for singlet cell events, and setting a gate to only include mRuby-BFP+ single-copy integrant cells. From this population, we derived median GFP and TA-splitHalo signal from 10k cell events. We employed a two sample Kolmogorov–Smirnov test to assess the significance of differences between the non-normally distributed populations in our data. The null hypothesis was that the signal from the knock-in population (+KI) is lower than the signal from the population with no knock-in (-KI). As an example, for the GFP(1-10) stable cell line, we can reject that null hypothesis with high confidence in the 488nm channel. For each cell line, we also calculated the signal to background ratio (SBR) in 488 nm and 646 nm channels, by dividing the median fluorescence intensity of the +KI cells by the median of –KI cells in the same cell line.

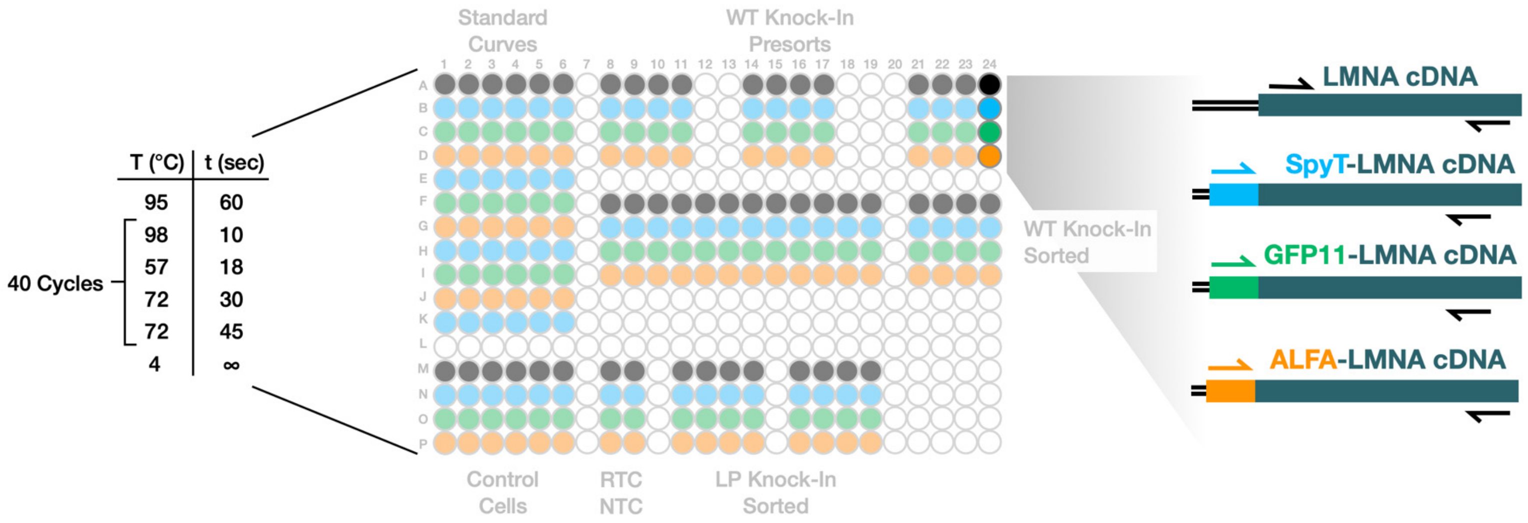


| Primer Set | Fwd Primer | Fwd Primer Sequence | Rev Primer | Rev Primer Sequence | Notes |
|---|---|---|---|---|---|
| Internal | LMNA_Fwd | CAGCTCCACTCCGCTGTC | LMNA_Rev | CGCTCCTTGGCTACTGAGTC | Amplifies all Lamin A/C mRNA |
| SpyT | SpyT-Fwd | CGCCTACAAGCGTTACAAGG | SpyT-Rev | CGCTCCTTGGCTACTGAGTC | Amplifies all SpyT-LMNA KIs |
| GFP11 | GFP11-Fwd | CGTGACCACATGGTCCTTC | GFP11-Rev | CTGACCACCTCTTCAGACTCG | Amplifies all GFP11-LMNA KIs |
| ALFA | ALFA-Fwd | GCCGACGATTGACTGAGC | ALFA-Rev | CTGACCACCTCTTCAGACTCG | Amplifies all ALFA-LMNA KIs |

**Figure S7:** qPCR layout is shown (top). Control and knock-in samples from all experiments are run using four internal (black), SpyT (blue), GFP11 (green), and ALFA (orange) primer sets on the same 384-well PCR plate. The internal primer set provides a readout of LMNA cDNA abundance which the KI specific sets can be compared to. Standard curves were produced using a 1:10 dilution series of plasmids containing the predicted amplicon.

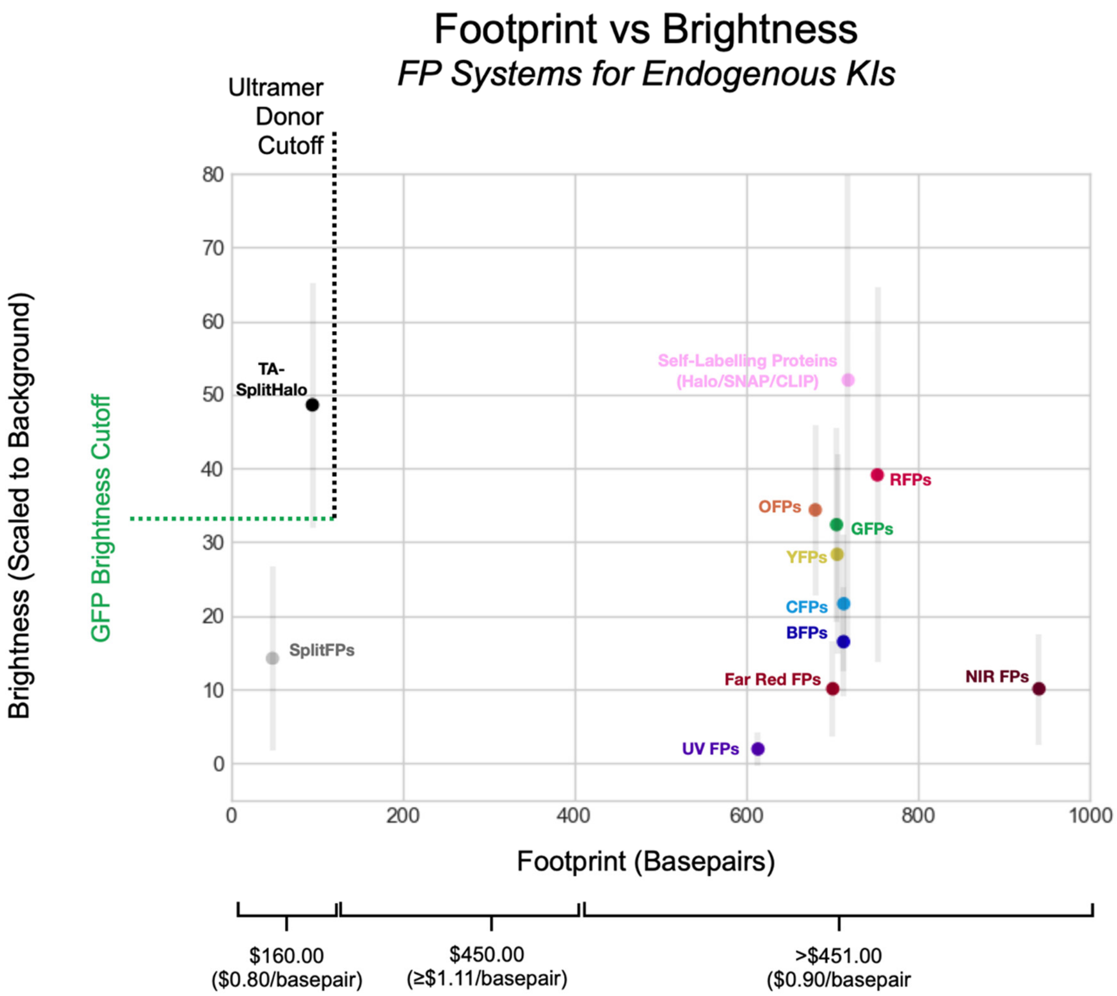


**Figure S8:** Scatterplot showing average brightness of fluorophores with respect to genomic footprint in base pairs and cost of producing ssDNA at given lengths. As shown, TA-splitHalo systems is the only platform that retains the cutoff for using ultramer size ssDNA donors while improving on the brightness of full-length fluorescent proteins on average.

| Primer Name | Primer Sequence | Notes |
|---|---|---|
| **ML557** | TAATACGACTCACTATAG | **Forward amplification primer.** |
| **ML558** | AAAAAAAGCACCGACTCGGTGC | **Reverse amplification primer.** |
| **LMNA-Specific Oligo** | TAATACGACTCACTATAG**CCATGGAGACCCCGTC CCAG**GTTTAAGAGCTATGCTGGAA | **Contains T7 transcription site LMNA-specific protospacer sequence.** |
| **ML611** | AAAAAAAGCACCGACTCGGTGCCACTTTTT CAAGTTGATAACGGACTAGCCTTATTTAAACTTGC TATGCTGTTTCCAGCATAGCTC TTAAAC | **Contains constant gRNA stem-loop.** |

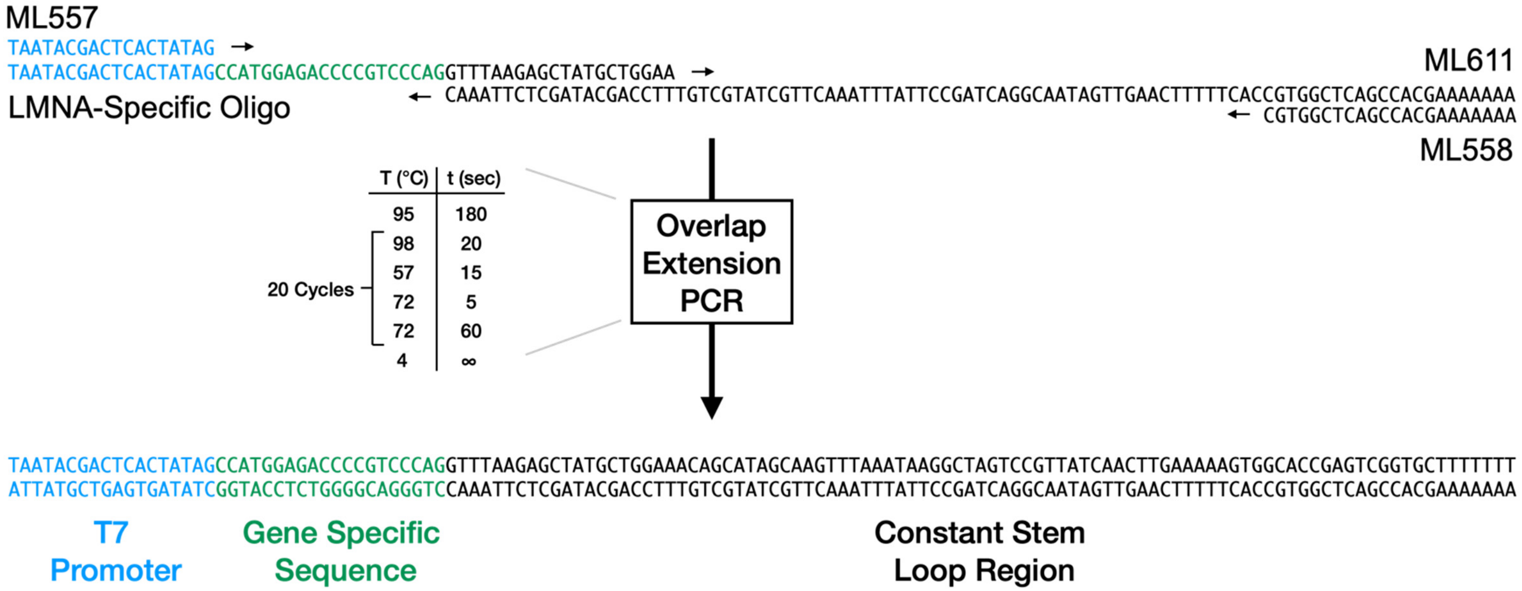


**Figure S9:** Methods detailing gRNA IVT template synthesis. Primer names, sequences, and contributions are shown (top) as is the PCR amplification scheme and final IVT template for LMNA gRNA generation containing the T7 promoter (blue), LMNA-specific PAM sequence (green), and constant stem-loop region (black) (bottom).

**Table S1:** 200 bp ultramer donor strands used for LMNA KIs. Sequences for GFP11 (green), SpyT (blue), ALFA (orange) GS linkers (black) and LMNA homology (teal) are bolded.

| Ultramer Name | Ultramer Sequence |
|---|---|
| GFP11-SpyT-LMNA | TTTCCGGGACCCCTGCCCCGCGGGCAGCGCTGCCAACCTGCCGGCCATG**CGTGACCACATGGTCCTTC**<br>**ATGAGTATGTAAATGCTGCTGGGATTACAGGTTCTGTGCCTACTATCGTGATGGTGGACGCCTACAA**<br>**GCGTTACAAGGGATCCGAGACCCCGTCCCAGCGGCGCCACCCGCAGCGGGGCGCAGGCCAGCT** |
| ALFA-SpyT-LMNA | CCTTTCCGGGACCCCTGCCCCGCGGGCAGCGCTGCCAACCTGCCGGCCATG**CCTAGCCGCCTGGAGGA**<br>**AGAACTCCGCCGACGATTGACTGAGCCAGGTTCTGTGCCTACTATCGTGATGGTGGACGCCTACAAG**<br>**CGTTACAAGGGATCCGAGACCCCGTCCCAGCGGCGCCACCCGCAGCGGGGCGCAGGCCAGCTC** |
| GFP11-LMNA | GTCCTTCGACCCGAGCCCCGCGCCCTTTCCGGGACCCCTGCCCCGCGGGCAGCGCTGCCAACCTGCCG<br>GCCATG**CGTGACCACATGGTCCTTCATGAGTATGTAAATGCTGCTGGGATTACAGGATCCGAGACCC**<br>**CGTCCCAGCGGCGCCACCCGCAGCGGGGCGCAGGCCAGCTCCACTCCGCTGTCGCCCACCCGC** |
| AFLA-LMNA | TGTCCTTCGACCCGAGCCCCGCGCCCTTTCCGGGACCCCTGCCCCGCGGGCAGCGCTGCCAACCTGCCG<br>GCCATG**CCTAGCCGCCTGGAGGAAGAACTCCGCCGACGATTGACTGAGCCAGGATCCGAGACCCCGT**<br>**CCCAGCGGCGCCACCCGCAGCGGGGCGCAGGCCAGCTCCACTCCGCTGTCGCCCACCCGCAT** |
| SpyT-LMNA | TCTGTCCTTCGACCCGAGCCCCGCGCCCTTTCCGGGACCCCTGCCCCGCGGGCAGCGCTGCCAACCTGC<br>CGGCCATG**GTGCCTACTATCGTGATGGTGGACGCCTACAAGCGTTACAAGGGATCCGAGACCCCGTC**<br>**CCAGCGGCGCCACCCGCAGCGGGGCGCAGGCCAGCTCCACTCCGCTGTCGCCCACCCGCATC** |
